# Mismatch type impacts interference and priming activities in the type I-E CRISPR-Cas system

**DOI:** 10.64898/2026.01.24.701482

**Authors:** Phong T. Phan, Meric Ozturk, Elizabeth M. Dougherty, Jerusha Ravishankar, Chaoyou Xue, Dipali G. Sashital

**Affiliations:** Roy J. Carver Department of Biochemistry, Biophysics and Molecular Biology, Iowa State University, Ames, IA 50011, USA

**Author notes:** Corresponding author: Dipali Sashital. Authors contributed equally (Phong T. Phan, Meric Ozturk). The American University in VietNam, 299 Tran Dai Nghia St, Hoa Hai Ward, Ngu Hanh Son District, Da Nang City, Vietnam (Phong T. Phan). State Key Laboratory of Engineering Biology for Low-Carbon Manufacturing, Tianjin Institute of Industrial Biotechnology, Chinese Academy of Sciences, Tianjin 300308, China (Chaoyou Xue).

## Abstract

Type I-E CRISPR-Cas systems direct RNA-guided interference against foreign nucleic acids using the CRISPR RNA (crRNA)-guided Cascade complex and Cas3 helicase-nuclease. DNA targeting by Cascade-Cas3 promotes priming, a mechanism that allows for rapid acquisition of new spacers within the CRISPR array. Target mutations in the PAM and PAM-proximal seed region can block interference but may still allow priming. Previous studies have suggested that target mutations to T and A are tolerated, but that C and G substitutions are deleterious to interference and priming, respectively. However, the contributions of the crRNA spacer sequence to mutational tolerance remain unclear. Here, we systematically tested the effects of crRNA seed sequences on mutational tolerance. We engineered four *E. coli* strains with variable spacer sequences and tested CRISPR interference and priming against a plasmid library for each strain. Consistent with prior studies, we observe that mutations to C or G in the seed can be highly deleterious, especially at positions 1, 2 and 4. However, the corresponding crRNA sequence also strongly impacts the level of defect, with rC-dC and rA/G-dG causing the largest defects in our plasmid library experiments. Using *in vitro* biochemistry, we observe that mismatch type at the first position of the seed affects Cas8 conformation, and results in reduction in the rates of both Cascade-target binding and Cas3 recruitment. Overall, our results reveal that although nucleotide identity of target mutations is an important determinant of type I-E CRISPR immunity, the crRNA sequence also strongly impacts immune outcomes upon target mutation.

## Introduction

CRISPR–Cas (clustered regularly interspaced short palindromic repeats – CRISPR associated) systems are RNA-guided immune systems that allow archaea and bacteria to defend against the constant threat from viruses and other invasive elements [1–4]. During initial infection, a short foreign DNA fragment, called a spacer, is inserted into the CRISPR array of the host chromosome via adaptation [1,5–7]. The CRISPR array is transcribed and the transcript is processed into mature CRISPR RNAs (crRNAs) [2,8–12], each carrying a unique spacer sequence that matches with the invader DNA and associates with Cas proteins to form an effector complex [2,13,14]. The effector complex uses the crRNA to bind to the complementary target region upon subsequent infection, leading to the destruction of foreign DNA via interference [2,3,13,15,16]. In some systems, spacers are preferentially acquired from DNA that is targeted by the Cas effector via priming, a positive feedback mechanism that enables rapid adaptation against an invader [17–22].

*E. coli K12* has a class 1, type I, subtype E (I-E) CRISPR–Cas immune system, which consists of two CRISPR loci and eight *cas* genes located in two operons (Fig. 1A) [23]. The crRNA effector complex, termed Cascade, comprises a crRNA and an unequal stoichiometry of five Cas proteins, Cas8 (also known as CasA or Cse1), Cas11 (CasB or Cse2), Cas7 (CasC or Cse4), Cas5 (CasD) and Cas6 (CasE or Cse3) [2,24]. In addition, Cas3, a protein containing both a nuclease and helicase domain, is required for target DNA degradation [2,25–29].

**Figure 1:**
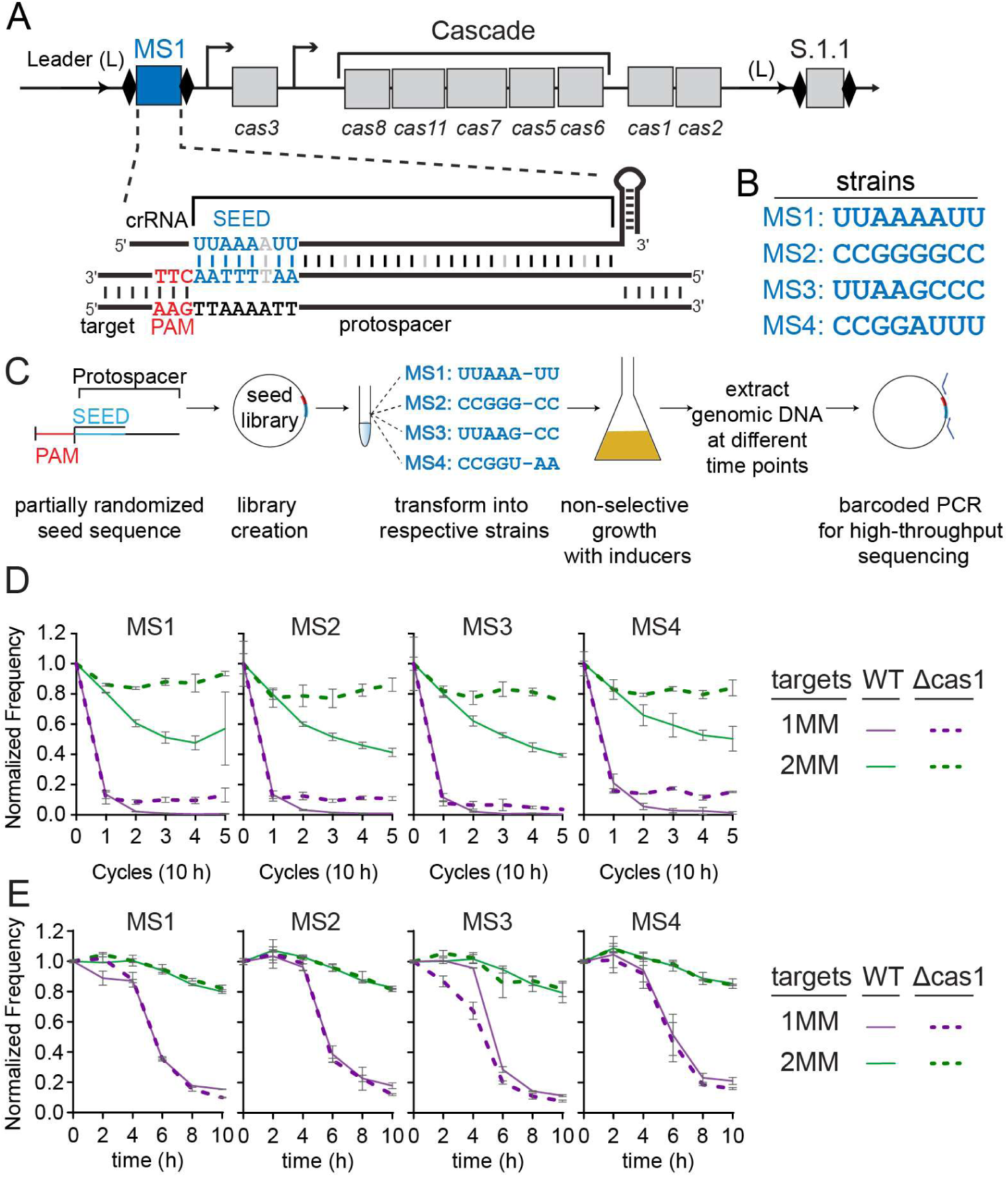
Timescales of direct interference and priming in an inducible type I-E CRISPR-Cas system. (A) Illustration of CRISPR-Cas operon from the modified *E. coli* strain used in the study. The strain contains two CRISPRs, each containing two repeats (black diamonds) and one spacer (rectangles labeled MS1 in CRISPR 2 and S.1.1 in CRISPR 1). The *cas* operon contains the *cas3* gene under IPTG-inducible *lac*UV5 promoter and the *cas8, cas11, cas7, cas5, cas6, cas1, cas2* operon under arabinose inducible *araBp8* promoter. Spacer sequences used in this study were introduced into the CRISPR 2 locus (shown in blue). A perfect seed target is shown for the MS1 sequence. (B) Seed sequences for spacer 2.1 for the four different strains. The remainder of the spacer sequence is identical. (C) Workflow of plasmid library selection assay. Four different partially randomized seed sequence libraries were created. Each library was then transformed to corresponding strains and grown for five 10-h cycles or for 10 hours with 2-h time points with induction of *cas* operon and without antibiotic selection. At each time point, the plasmid library was extracted and the target region was PCR amplified and submitted for high-throughput sequencing. (D-E) Overview of seed library sequence selection for WT and Δ*cas1* strains over five 10-hour cycles (D) and 2-h time points (E). Normalized frequency of targets containing one mismatch (1MM, purple) and targets containing two mismatches (2MM, green) over time for WT (solid line) and Δ*cas1* (dashed line). Fraction of reads for all 1MM or 2MM sequences in each strain were first normalized to the 3-5MM reads (see Methods) and then to the fraction of reads present at cycle or time point 0. The average of two replicates is shown, and error bars are the standard deviation with error propagated for the normalization.

During infection, Cascade locates target sequences in the invader DNA by initially searching for the PAM (protospacer adjacent motif) sequence [30,31], which is recognized by the Cas8 subunit of Cascade [21,32,33]. PAM recognition triggers DNA unwinding and the formation of an RNA-DNA hybrid duplex between the guide crRNA and the DNA target [33,34]. Target unwinding occurs directionally away from the PAM [35], and base pairing of the first 8 nucleotides with the guide RNA, called the “seed sequence”, is extremely important to ensure binding between the crRNA and the target DNA [19,20,31]. Mutations within the PAM and/or seed region block Cascade binding and lead to the escape from CRISPR interference, but may still enable primed spacer acquisition, resulting in the acquisition of new spacers that drive interference [17,18].

Although seed mutations are generally detrimental to CRISPR interference, interference efficiency varies depending on the type of mutation, and location within the seed [19,20,36,37]. Previous studies have found that target mutations to G and C are more deleterious than mutations to A and T, and that mutations at position 4 of the seed can be as deleterious as mutations at the first two positions of the seed [20,37,38]. While these mutations are generally thought to reduce the rate of Cascade-target binding [31,38], mutations within the seed sequence also affect Cascade conformation, particularly to the Cas8 subunit [39,40]. Because Cas8 directly interacts with Cas3 during interference [27,29], alternative conformations of Cas8 are thought to negatively impact Cas3 recruitment [29,33,39,41,42]. Thus, Cas8 conformation, in addition to defects in target binding, may contribute to the ability of Cascade to direct interference against a target.

A previous study suggested that crRNA spacer sequence can impact interference efficiency, with increased guanine/cytosine (G/C) content providing higher efficiency [43]. The spacer sequence may also impact mutational tolerance in the seed, as different mismatch types would be introduced depending on the crRNA sequence [19]. However, studies directly comparing the impact of mismatch type within the seed are limited. To address this limitation, we engineered four different *E. coli* strains expressing varied crRNA seed sequences with otherwise identical spacer sequences. Using these strains, we systematically tested the functionality of G/C rich or poor seed sequences using high throughput screening of a target mismatch library. Consistent with previous studies, we show that C or G mismatches at specific positions within the seed are more deleterious than others, although the corresponding ribonucleotide in the crRNA can impact the level of defect. Using complementary in vitro approaches, we observe that C and G mutations impact Cas8 conformation more substantially than T mutations, leading to reduced rates of target degradation when Cascade is pre-bound to the target DNA. Overall, these results indicate that type I-E immune activities are impacted not only by the mutation type in the target, but also by the mismatch with the crRNA, and can result in decreased kinetics of both target binding and Cas3 recruitment.

## Results

### Evaluating the effects of 1 or 2 mismatches for seed sequences with different G/C contents

To systematically evaluate how different mismatch types within seed sequence influences CRISPR interference and priming against mismatched targets, we created four strains of *E. coli K-12* in which the G/C content of the seed sequence of one spacer was varied, while the remainder of the spacer sequence remained constant. The strains were derived from a modified BW25113 strain [44], in which the *cas* genes are under control of inducible promoters (Fig. 1A). Each strain expresses two crRNAs, one from each of the two CRISPRs in the *E. coli* genome. The four seed variants were designed as follows: MS1, A/U-rich; MS2, G/C-rich; MS3, A/U in the first four positions and G/C in the last four; and MS4, G/C in the first four positions and A/U in the last four (Fig. 1B). For each strain, a Δ*cas1* deletion was also engineered. Because the Δ*cas1* strain cannot acquire new spacers, comparison of target loss in Δ*cas1* and wild-type strains allows for the differentiation of target loss through direct interference by the original spacer or target loss through the acquisition of new spacers via priming [7,19].

For each strain, we carried out an *in vivo* plasmid library selection assay followed by high-throughput sequencing (HTS). In this approach, target libraries containing seed sequence mismatches were constructed for each strain, with the majority of sequences containing one or two mismatches (Fig. 1C, S1A). The sixth position of the crRNA does not form a base-pair with the target DNA, so was not varied within the library (see Methods) [31,45,46]. Each library was introduced into the corresponding strain and subjected to negative selection by CRISPR immunity over five consecutive 10-hour growth cycles (Fig. 1C). After each cycle, the target region of the plasmid library was PCR-amplified and subjected to HTS. Sequences that progressively disappear represent mutations that remain susceptible to either direct interference (i.e. lost in both Δ*cas1* and WT) by the original spacer or to priming (i.e. lost in WT but not in Δ*cas1*).

Although the plasmid library was designed to strongly represent sequences containing one or two mismatches between crRNA and target, it also contained targets with three or more mismatches. As expected from previous studies, we found no evidence of direct interference against sequences carrying three or more mismatches in the seed sequence [19,20], as indicated by their relative enrichment throughout the cycles (Fig. S1A). Because these mismatched targets should remain stable over time, we normalized our datasets so that the proportion of sequences with three to five mismatches remained constant (see Methods) [47]. This approach more accurately reflects the depletion of sequences with one or two mismatches from the library (Fig. S1B).

The normalized frequency of sequences having one or two mismatches (1MM and 2MM) revealed that 1MM targets were rapidly depleted by the end of cycle 1 in both WT and *Δcas1* strains (Fig. 1D). A small fraction of 1MM sequences remained stable in the Δ*cas1* strain, as indicated by their slightly increased frequencies at later time points relative to the WT strain. Notably, 1MM sequences were depleted more in the MS3 *Δcas1* strains relative to the other three Δ*cas1* strains, suggesting that the MS3 crRNA is more tolerant of mutations for direct interference. The normalized frequency of 2MM sequences declined over time in all WT strains but remained enriched in the *Δcas1* strains, indicating that 2MM sequences mostly block direct interference in Δ*cas1* but can support priming in WT strains, similar to previous studies (Fig. 1D) [19,20].

Most of the 1MM sequences were depleted within the first 10-h growth cycle, limiting our ability to analyze the effects of each mismatch type. We therefore performed the same plasmid library depletion assay using 2-h time points within the timeframe of the first 10-h growth cycle. We did not see any significant difference in the loss of 1MM or 2MM sequences between the WT and Δ*cas1* strains across these shorter time points (Fig. 1E), consistent with the similar frequency of sequences between the two strains at cycle 1 (Fig. 1D). Similarly, when we assessed spacer acquisition through amplification of the CRISPR 2 array for each WT strain bearing the target plasmid library, we observed expanded CRISPRs only following the first 10-h cycle (Fig. S2). Overall, these results indicate that sequences are lost only by direct interference through the first 10 hours following induction in our engineered strain, and that target loss through priming begins to occur only in the second 10-h cycle.

### Effects of individual mismatch type and position on CRISPR immunity

As observed in Fig. 1E, 1MM sequences decayed exponentially following a delay in both the WT and Δ*cas1* strains over a 10-h time period. Most individual 1MM sequences also displayed a similar delayed decay (Fig. S3). We fit the normalized frequency of individual 1MM sequences over the 2-h time points to a model that accounts for this time delay prior to exponential delay, and calculated half-lives for each sequence (Fig. 2A, S3, see Methods). Because the 1MM sequences decayed at similar rates in WT and Δ*cas1* strains (Fig. 1E), we included the half-life values for both sets of strains in our analysis.

**Figure 2:**
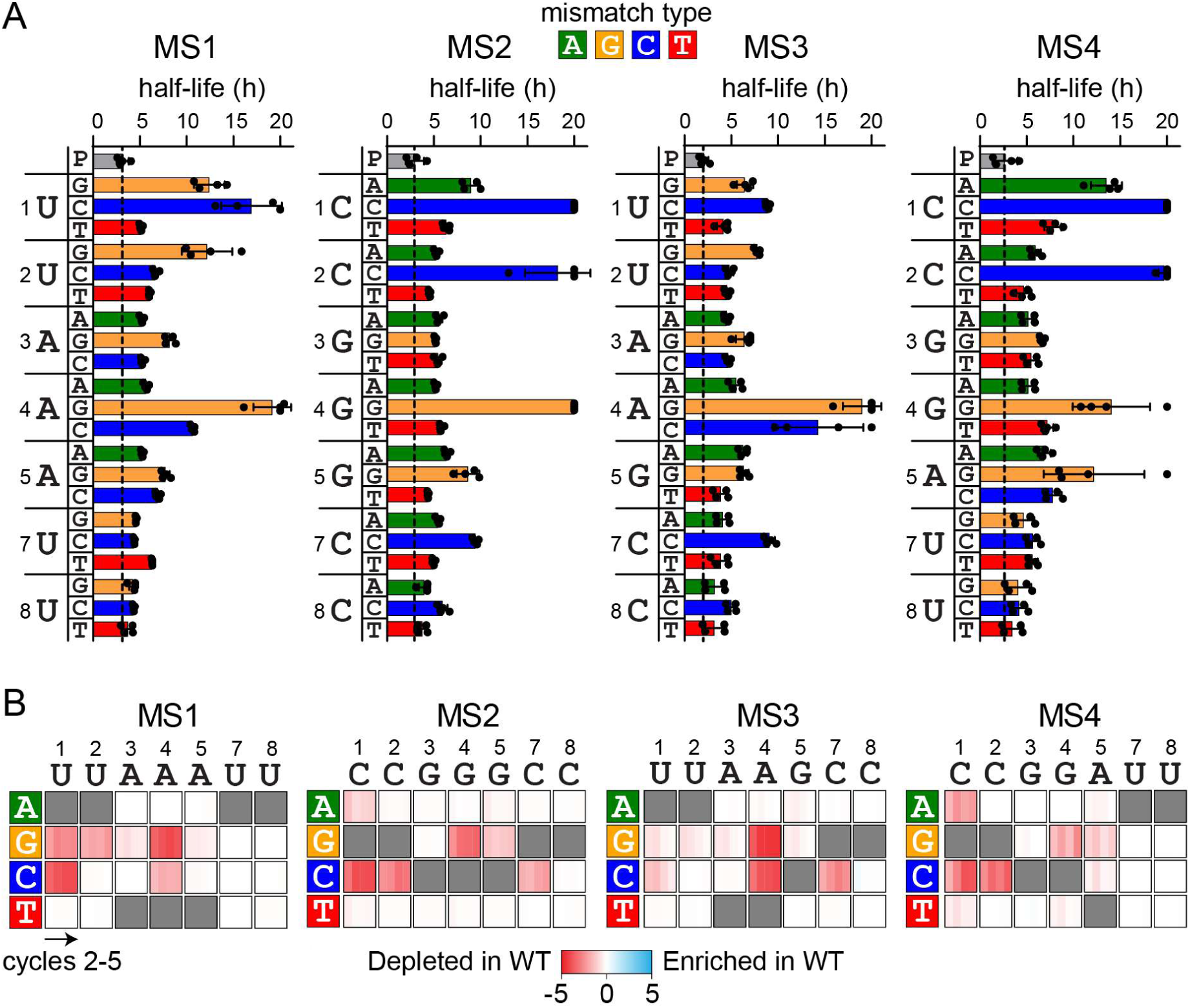
Influence of mismatch type and location on direct interference and priming against 1MM seed mutants. (A) Half-lives for sequences containing a single mismatch at each position of the seed for MS1-MS4. Dashed line represents the half-life for the perfect target (P, gray bar). The crRNA sequence is listed to the left of the axis, and half-lives for the three potential mismatches are plotted at each position measured from the short time-point data (see Fig. S3 for decay curves). Mismatch types are colored as follows: dA = green, dG = orange, dC = blue, dT = red. Half-lives were measured from two replicates from each strain, WT and Δ*cas1*. The bars represent the mean of these half-lives. Each data point is plotted as a black circle, with error bars representing the standard deviation. Sequences from individual replicates that did not decay sufficiently to fit to the delayed decay model were capped at a half-life of 20 h. (B) Heatmaps showing the depletion score for all 1MM target sequences in WT relative to Δ*cas1* strains for MS1-MS4 over 10-hour cycles. Low depletion scores (red) indicate that sequences were less abundant in WT than in Δ*cas1.* Depletion scores of 0 (white) indicate that sequences were equally abundant in WT and Δ*cas1.* Three possible nucleotide substitutions are plotted vertically (labeled on left) for each position of the crRNA (horizontal, labeled on top). Depletion scores for cycles 2-5 are plotted in each box. Gray boxes are shown for complementary target nucleotides. Values represent the average of 2 replicates.

**Figure 3:**
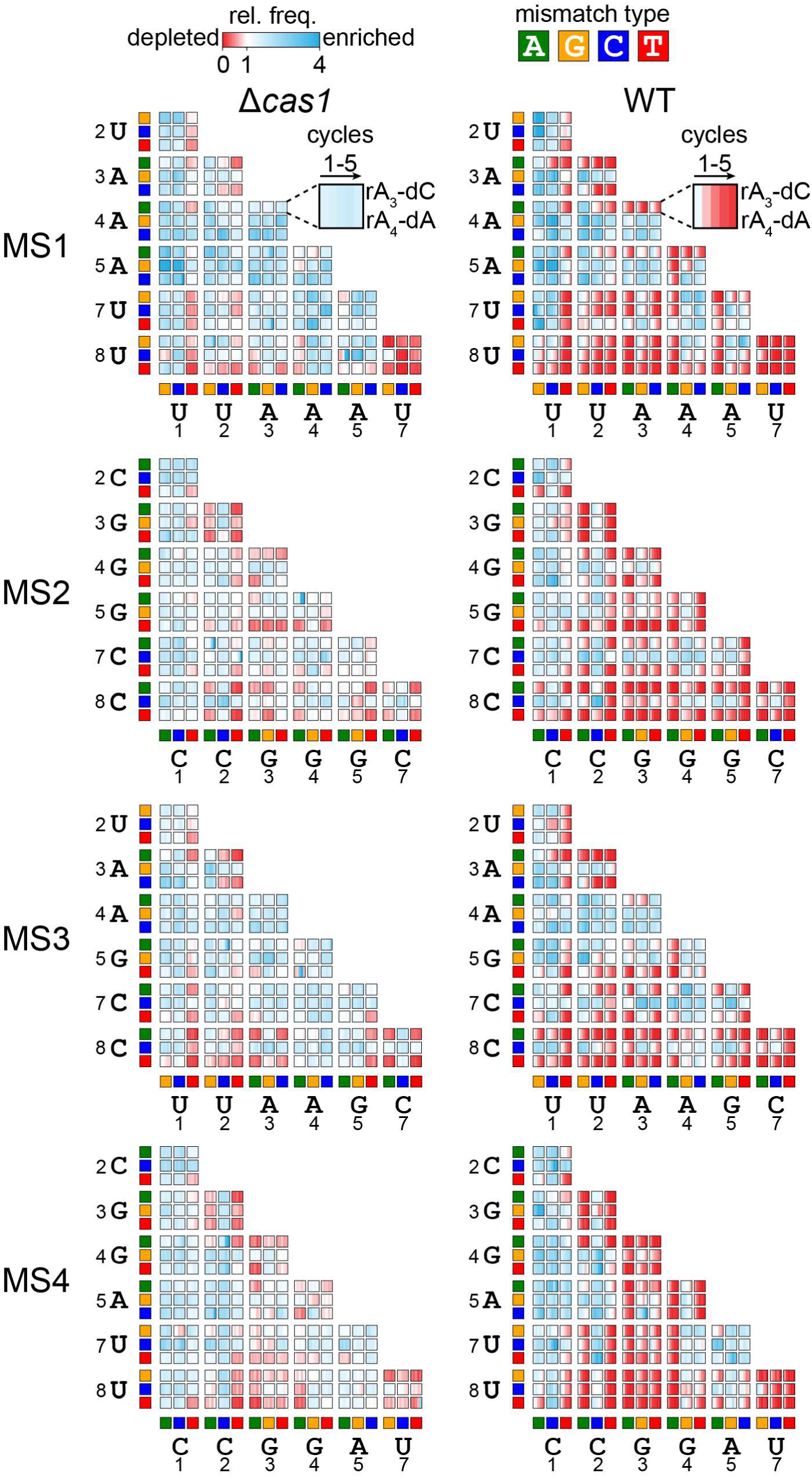
Comparison of two mismatch target depletion in WT and Δ*cas1* strains. Heatmaps showing the relative enrichment (blue) and depletion (red) of all possible 2MM target sequences over 5 cycles for all the Δ*cas1* (left) and WT (right) strains. The relative frequency (rel. freq.) of each 2MM sequence is plotted for the 10-h cycles 1-5 (see Methods). The relative frequency values represent the average of n = 2. Each 3 × 3 array represents the 9 possible mismatches at a given set of two positions. The perfect target sequences are not shown in the heatmap. The mismatch type is represented by colored boxes on the x-axis (positions 1-7) and y-axis (positions 2-8), with green representing dA, orange representing dG, blue representing dC, and red representing dT. An exemplar cell is highlighted for the MS1 Δ*cas1* and WT samples, showing the combination of rA-dC at position 3 of the crRNA and rA-dA at position 4 of the crRNA.

Mismatches at positions 1, 2 and 4 of the seed sequence containing C or G in the target sequence caused the longest half-lives, similar to prior studies [19,20,38]. However, the severity of the effect varied by mismatch type. In particular, sequences that introduced rC-dC mismatches at positions 1 and 2 of MS2 and MS4 were stable and did not decay over the 10-h time period (Fig. S3), while rC-dC also caused the longest half-lives at position 7. At position 4 of MS1 and MS3, rA-dC was also relatively deleterious. In contrast, while an rU-dC mismatch was deleterious at position 1 of MS1, rU-dC was relatively tolerated at position 2 of MS1 and at both positions 1 and 2 of MS3 (Fig. 2A). For mutations to dG, both rA-dG (MS1 and MS3) and rG-dG (MS2 and MS4) mismatches at position 4 were more deleterious than rU-dG at positions 1 and 2 (MS1 and MS3). While rC-dA at position 1 of MS2 and MS4 caused relatively slow decay, rC-dA was tolerated at position 2, as was either rA-dA or rG-dA at position 4 of all four strains. Mismatches involving dT were generally tolerated at all positions regardless of the RNA sequence, with rC-dT (MS2 and MS4) causing a slight increase in half-life in comparison to rU-dT (MS1 and MS3) at position 1. Notably, although the four strains contain differing neighboring nucleotides at positions 4 and 5, we observed relatively little difference between the same mismatches at these positions (Fig. 2A), suggesting that nearest neighbor effects do not substantially influence mutational tolerance within the seed. However, the differences observed at the first two positions of MS1 and MS3 do suggest that overall sequence context may impact mutational tolerance to some degree. Overall, our results indicate that mismatch type between the crRNA and target DNA has a substantial impact on mismatch tolerance within the seed sequence.

We next compared how 1MM sequences were depleted in WT versus *Δcas1* strains over the longer, 10-h cycles. As described above, target loss through priming occurs following the first 10-h cycle (Fig. 1D, S2), and is expected to result in the loss of 1MM sequences that were stable over the first 10-h cycle (Fig. 2A, long half-lives). We calculated depletion scores that measure how the abundance of each 1MM sequence diverges between WT and Δ*cas1* (see Methods). Sequences with half-lives shorter than 10 hours (Fig. 2A) had depletion scores near 0 (Fig. 2B), indicating that they were similarly depleted via direct interference in both WT and Δ*cas1* by the end of the first cycle. Sequences with longer half-lives (Fig. 2A) had larger negative depletion scores (Fig. 2B), indicating that they were depleted in WT but remained abundant in Δ*cas1.* These sequences cannot undergo direct interference but can promote priming, resulting in the loss of these sequences starting in cycle 2 in the WT strains. In general, all 1MM sequences that caused defects in direct interference over the first 10-h cycle promoted priming, with the strongest priming-dependent target loss observed for sequences with the largest direct interference defects (compare Fig. 2B and 2A).

### Combinations of mismatches in the seed region promote priming

We next examined all target sequences containing two seed mismatches (2MM) in Δ*cas1* and WT strains over time (Fig. 3). As predicted from the overall library analysis (Fig. 1D–E), only a small number of 2MM sequences were depleted in *Δcas1* strains, whereas many were progressively depleted in WT, showing that these sequences enable priming. The *Δcas1* heat maps highlight which mismatch combinations can still support direct interference. Sequences containing one mismatch at position 8 in combination with a second mismatch could often undergo direct interference. This result is expected, as mismatches further away from the PAM become progressively less deleterious [20,38]. However, a rC-dC pair at position 8 of MS2 and MS3 is deleterious in combination with most other mismatches, while several sequences containing rU-dC at position 8 of MS1 and MS3 were depleted in the Δ*cas1* strain, underscoring the relative severity of rC-dC mismatches versus rU-dC. Conversely, rU-dT at position 1 of MS1 and MS3 is relatively well tolerated in comparison to rC-dT at the same position of MS2 and MS4. Surprisingly, some combinations of mismatches at adjacent positions were tolerated. For example, sequences with combinations of rC-dA or rC-dT at position 2 and any mismatch at position 3 were relatively depleted in MS2 and MS4, while sequences with combinations of rU-dC and rU-dT at position 2 and rA-dA and rA-dC at position 3 were relatively depleted in MS1 and MS3. Similar tolerance of adjacent mismatches was also observed for some mismatch combinations at positions 3 and 4 for MS2 and MS4, although rG-dG at position 4 resulted in complete loss of direct interference.

Comparison of the 2MM heatmaps for Δ*cas1* and WT strains indicates that many sequences that were stable in Δ*cas1* were lost via priming in WT. However, many sequences also remained stable in the WT strain, indicating sequences that are likely not bound by Cascade. Similar to the 1MM sequences that block direct interference (Fig. 2A), the 2MM sequences that block priming are dependent not only on nucleotide identity of the target mutation, but also on the mismatch type based on the crRNA sequence. For example, rA-dG at position 3 blocks priming for MS1 and MS3 when combined with mismatches at positions 1 and 2. In contrast, rG-dG at position 3 can be tolerated for MS2 and MS4, especially in combination with rC-dT at position 2. Nevertheless, for MS4, we did observe tolerance of rG-dG mismatches at position 4 and most mismatches at position 3, with the exception of a second rG-dG at this position. Thus, while both purine-purine mismatch types involving dG may be detrimental to Cascade binding, they can be tolerated in some combinations allowing priming to still occur.

Overall, the combined analysis of 1MM and 2MM sequences that can undergo direct interference and priming, respectively, reveal the importance of the crRNA-DNA mismatch type in dictating the severity of defects for dG and dC target mutations. While both purine-purine mismatches involving dG can be more deleterious than rU-dG, rC-dC is substantially more deleterious than rU-dC or rA-dC.

### Mismatch type affects Cas8 conformation and target binding dynamics

We next investigated how different mismatch types impact Cascade conformation and Cas3-mediated degradation rates. Previous work has shown that Cas8 adopts distinct conformations depending on the type of target bound by Cascade [39]. The Cas8 N-terminal domain can adopt either a closed or an open conformation, while the C-terminal four-helix bundle can adopt either an unlocked or a locked conformation [33,45,48,49]. Cascade adopts a closed and unlocked conformation when unbound to a perfectly matched double-stranded DNA target [48,49]. The closed and locked conformation exposes Cas8 residues that contact Cas3, facilitating Cas3 binding and recruitment to the target DNA [29,33,41,42]. Cas8 shifts to the open conformation upon single-stranded DNA binding or when mutations are present in the PAM or the seed of double-stranded DNA targets [39,45], a conformation that is thought to block Cas3 recruitment [29,41].

To examine how different mismatch types influence Cas8 conformation, we performed a previously developed fluorescence resonance energy transfer (FRET) assay [39] using Cascade bearing the MS1 crRNA. In this assay, the N-terminal domain or C-terminal domain of Cas8 is labeled with Cy3 and the Cas5 subunit is labeled with Cy5 (Fig. 4 A, B). The N-terminal domain of Cas8 moves towards Cas5 when it adopts open conformation, causing an increase in FRET signal (Fig. 4A). The C-terminal domain of Cas8 moves towards Cas5 when it adopts locked conformation (Fig. 4B).

**Figure 4:**
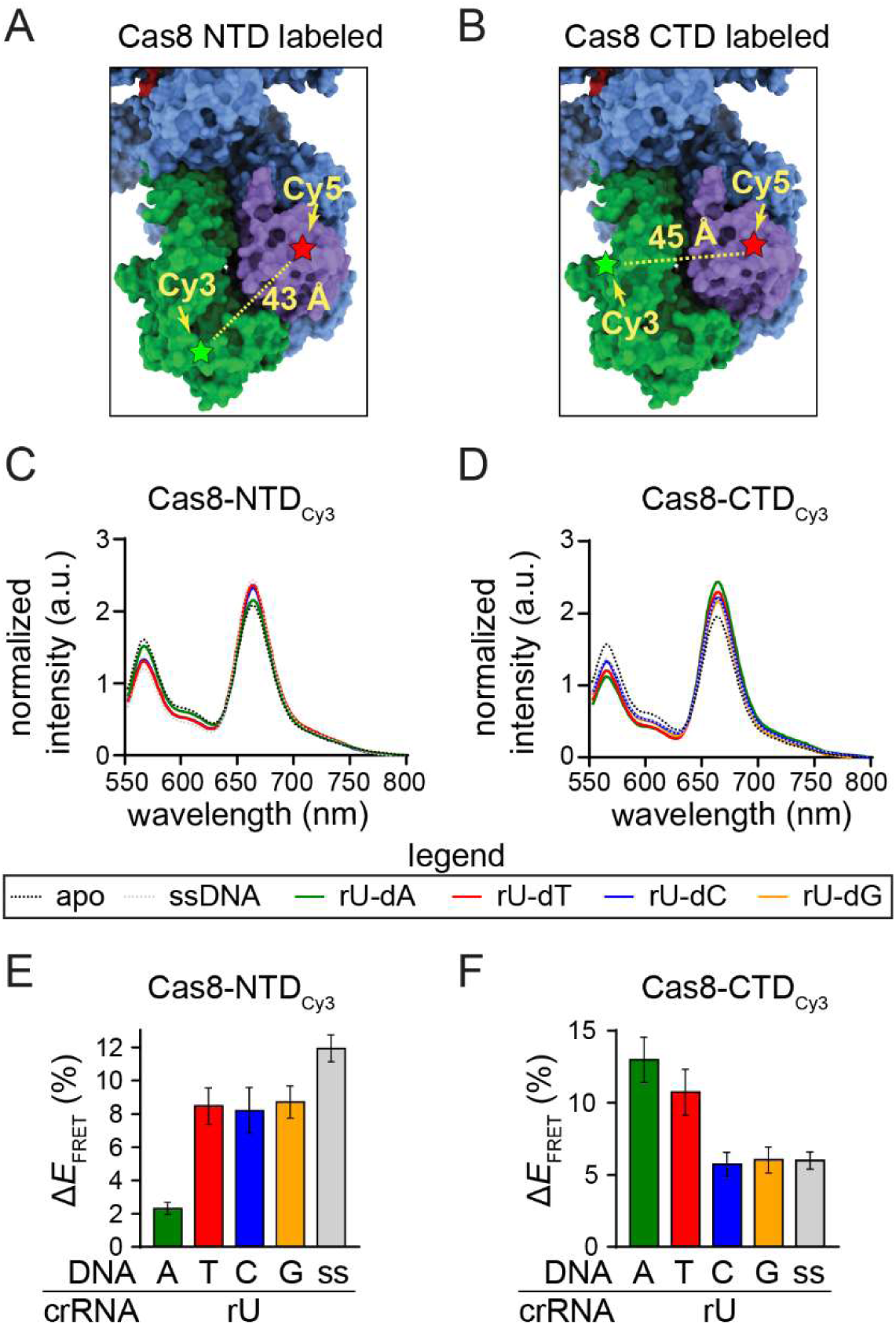
Measuring Cas8 conformation dependence on mismatch type using FRET. The position of the Cy3 label (A) on the N-terminal domain of Cas8 (Cys 69) or (B) on the C-terminal domain of Cas8 (Cys 376); Cy5 is on Cas5 (Cys 169). The estimated distance between dyes is based on distance between C-alpha of the two labeled residues in the apo Cascade crystal structure (PDB ID 4TVX). (C, D) Normalized FRET intensity for NTD- and CTD-labeled Cascade bound to single stranded DNA (ss) or double-stranded perfect target (rU-dA), or targets introducing an rU-dT, rU-dC, or rU-dG mismatch at the first position of the target. (E, F) FRET efficiency changes for NTD- and CTD-labeled Cascade bound to different targets. The change in FRET was measured relative to apo Cascade that was not bound to DNA (see Methods). The average of three replicates is shown, with error bars representing the propagated standard deviations. Individual graphs and controls are shown in Fig. S4.

We measured Cas8 conformation when Cascade is bound to double-stranded DNA (dsDNA) targets containing either a perfectly matched DNA (rU-dA at position 1) or targets containing all three mismatches at position 1. A single-stranded (ss) DNA target was also included as a positive control for the open conformation. When the N-terminal domain of Cas8 was labeled, all three mismatched targets had similar FRET efficiencies relative to the *apo* Cascade control (Fig. 4 C, E, S4A). FRET efficiencies for the mismatched targets were between the perfect dsDNA and ssDNA controls, suggesting that the three mismatched targets induce the open conformation to a similar degree that exceeds the perfect dsDNA target, but not to the same degree as the ssDNA target. In contrast, when the C-terminal domain of Cas8 was labeled, we observed similar FRET efficiencies relative to *apo* Cascade for the perfect target and the rU-dT mismatch, while rU-dC and rU-dG had FRET efficiencies more similar to the ssDNA control (Fig. 4 D, F, S4B). These findings suggest that Cas8 adopts an alternative conformational ensemble when MS1-Cascade binds to the dT mismatched target, relative to the dC and dG targets.

To determine whether the conformational differences induced by the dC and dG mismatched target impacts Cas3-dependent target degradation, we performed Cascade-Cas3 cleavage assays against plasmid targets bearing either the perfect target or the three mismatched targets at position 1 (Fig. S5). In a simplified kinetic model, the overall rate of target degradation is based on the rate of target binding by Cascade (*k*_bind_), Cas3 recruitment to Cascade (*k*_recruit_), and target cleavage by Cas3 (*k*_cleave_) (Fig. 5A). Mismatches at position 1 are expected to decrease the rate of target binding (*k*_bind_), which may mask the effects of Cas8 conformation on the rate of Cas3 recruitment and subsequent steps (*k*_recruit_ and *k*_cleave_). We therefore performed cleavage assays in two different ways. In the first assay, Cascade and Cas3 were pre-mixed prior to the addition of the DNA target, resulting in cleavage rates that are dependent on all three phases of our kinetic model (Fig. 5B). In the second assay, Cascade and the target were incubated together, allowing Cascade to pre-bind the DNA prior to addition of Cas3 (Fig. 5C). This experimental design eliminates the rate of Cascade-target binding from the kinetic model and should more directly inform on the rate at which Cas8 recruits Cas3.

**Figure 5:**
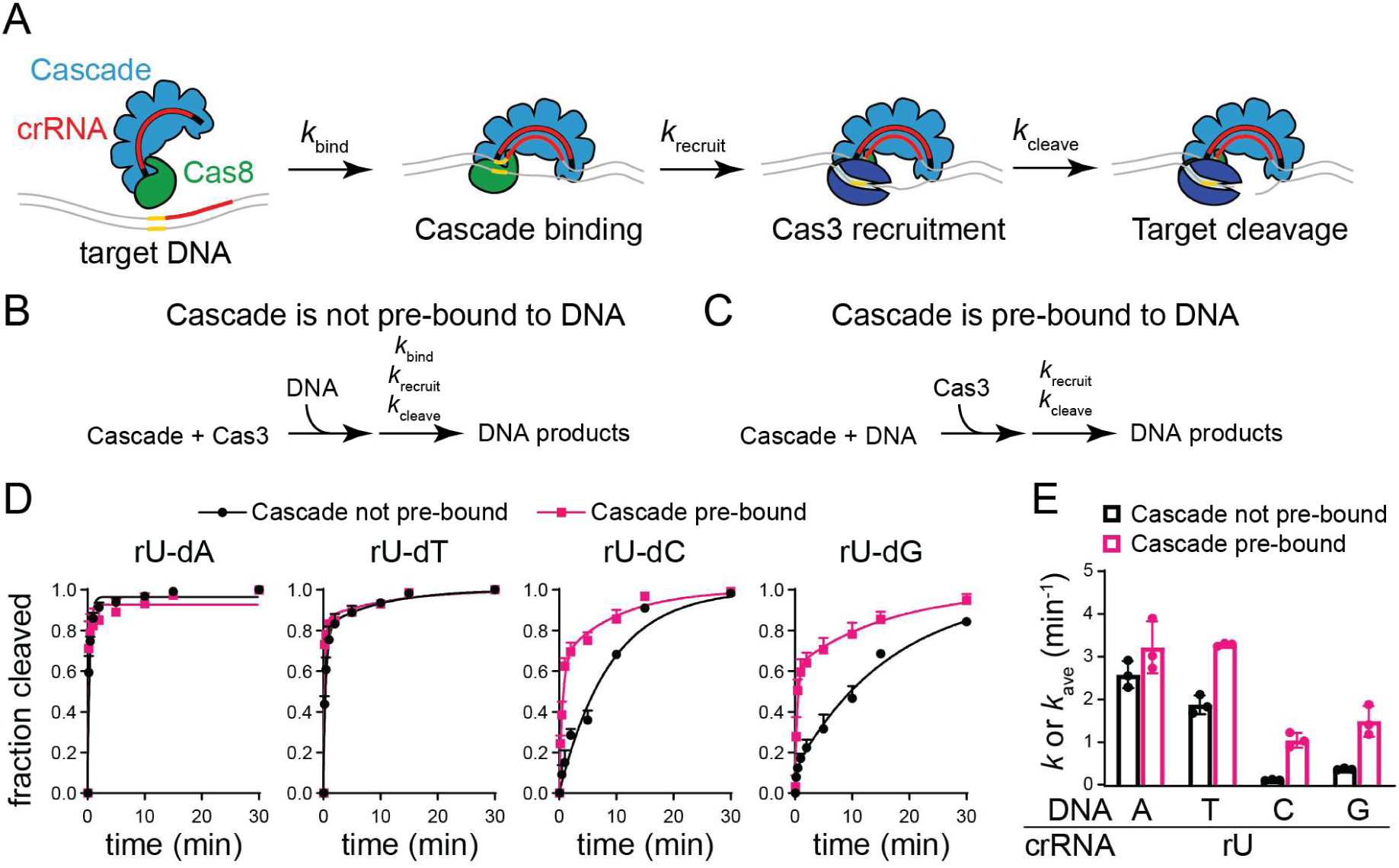
Measuring the effects of mismatch type on the rate of target binding and degradation. (A) Kinetic steps that determine the rate of target degradation by Cascade-Cas3 complex. The rates of target binding (represented by *k*_bind_), Cas3 recruitment (*k*_recruit_), and target cleavage (*k*_cleave_) all contribute to the overall rate of target degradation. (B, C) Schematic view of experimental design. (B) When Cascade and Cas3 are mixed together prior to addition of DNA, target degradation is dependent on the rate of target binding, Cas3 recruitment, and target cleavage. (C) When Cascade is incubated with DNA to allow pre-binding prior to addition of Cas3, the rate of target degradation is dependent only on the rate of Cas3 recruitment and target cleavage. (D) Fraction cleaved over time for Cascade is not pre-bound to DNA (black circles, experimental design shown in panel B) and Cascade is pre-bound to DNA (pink squares, experimental design shown in panel C). Data points are the average of three replicates and standard deviation is shown as error bars. The data were fit to a double-exponential rate equation, with the exception of the perfect target (rU-dA) in both conditions and the rU-dC target when Cascade was not pre-bound, which fit better to a single-exponential rate equation (see Methods). (E) Observed rate constants (*k*) or average observed rate constant (*k*_avg_) values for Cascade bound to different target. For targets that fit best to a double-exponential, *k*_avg_ was determined by averaging the fitted rate constants for the fast and slow phase, with the exception of the rU-dA target in both conditions and the rU-dC target when Cascade was not pre-bound, for which a single *k* value was determined (see Methods). The average of three replicates are shown, with individual data points represented with circles and error bars representing standard deviation.

We observed that the target degradation reactions were biphasic for most targets and fit well to a double-exponential rate equation, with the exception of the perfect (rU-dA) target in both assays and the rU-dC target when Cascade was not pre-bound to the DNA (Fig. 5D). To compare *k* values, we averaged the rate constants measured for the slow and fast phases for cleavage data that was fit to a double-exponential model to account for the biphasic nature of the reaction (see Methods) (Fig. 5E). We observed similar rates for the rU-dA and rU-dT targets, both of which were degraded much more rapidly than targets that introduce an rU-dC or rU-dG mismatch at position 1 for both types of assays (Fig. 5D). When Cascade was not pre-bound to the target, the rU-dT mismatch causes a slight decrease in the rate of cleavage (∼1.4x lower than the perfect target, p = 0.03 based on two-tailed, unpaired t-test). However, when Cascade was pre-bound to the target, we observed no significant difference in the rate of cleavage for the perfect target and the rU-dT mismatched target. These results suggest that the rU-dT target has a small effect on the rate of target binding, but that Cas3 recruitment and cleavage activity is unaffected by the differences in Cas8 conformation induced by the rU-dT mismatch at position 1.

For the other two mismatched targets, when Cascade was pre-bound to the DNA, we observed a 2-3x decrease in the rate of target degradation compared to the perfect target and the rU-dT mismatched target (Fig. 5E). When Cascade was not pre-bound to the DNA, this difference was substantially larger, with the rU-dC target causing a 23x decrease and the rU-dG target causing a 7x decrease compared to the perfect target. Collectively, these results indicate that both the rU-dC and rU-dG mismatches decrease the rate of Cas3 recruitment to a similar degree, while the rU-dC causes a larger defect in the rate of target binding in comparison to the rU-dG mismatch.

## Discussion

In this study, we have systematically investigated the effect of the crRNA seed sequence on CRISPR interference and priming in the type I-E CRISPR-Cas immune system. Our results confirm that depending on the position and type of mismatches, CRISPR immunity can tolerate up to two mismatches in the seed region and promote priming. Similar to previous studies, we observed that dC and dG mutations in the target strongly inhibit direct interference [20,37,38], especially when located at positions 1, 2 and 4 of the seed. However, our findings also implicate the crRNA nucleotide in mutational tolerance. For dC mutations, rU and rA in the crRNA provided greater tolerance than rC, while for dG mutations, rU provided greater tolerance than rG or rA.

Previous structural modeling of multiple mismatch types provided insight into the impact of the crRNA-DNA mismatch type on Cascade-target binding [37]. In particular, rA-dG creates a clash due to the bulkiness of purine-purine base pairing, while rU-dG creates a clash with the thumb region of the Cas7 backbone subunit. Although dC mismatches were better accommodated within these structural models, we note that rC-dC is among the most deleterious mismatch type in our plasmid loss experiments. Similarly, we observed that an rU-dC mismatch causes a larger binding defect than an rU-dG mismatch for the MS1 crRNA (Fig. 5E), consistent with the rate of direct interference that we observed in cells (Fig. 2A). Thus, it is unclear whether Cascade accommodation of mismatched target sequence alone is sufficient to explain the effects of certain types of mismatches. It is tempting to speculate that dC and dG mismatches are more deleterious to DNA unwinding by Cascade. However, while it is important to consider the overall energy of target unwinding, it is also essential to consider how the mismatch type that is introduced affects Cascade-target binding. Certain mismatches could potentially enable formation of a more stable R-loop than other mismatch types, which is required for Cas3 recruitment [29,35,41].

An additional consideration is the effect of mismatch type on Cascade conformation. The conformation of the Cas8 subunit is sensitive to mutations in or near the PAM [39,40], and the conformation of this subunit is also important for ensuring efficient Cas3 recruitment and target degradation [29,33,41,42]. We find that the rU-dT mismatch that is best tolerated for direct interference at position 1 of the MS1 target (Fig. 2A) adopts a different Cas8 conformation than rU-dC or rU-dC (Fig. 4F), resulting in more rapid cleavage of the target when Cascade is pre-bound to the target DNA (Fig. 5E). These collective results strongly suggest that mismatch type may impact the rate of interference not only by impacting the rate of target binding, but also by reducing the rate of Cas3 recruitment following target binding.

Overall, our study provides initial insight into the impact of mismatch type on type I-E interference and priming activities. An important limitation of our and previous studies is the difficulty of investigating all possible mismatch types at all positions [19,20,36,38]. Although our dataset covered four distinct crRNA sequences, it still lacks several mismatch types (e.g. rG or rA mismatches at the first two positions of the seed). Future studies may focus on the impact of other mismatch types, as well as the importance of mismatch type in the context of phage escape.

## Material and Methods

### Bacterial strains

Primers for *E. coli* engineering are listed in Table S1 and strains used in the study are listed in Table S2. All strains used in this study originated from BW25113 strain [44,50] which originally contained 18 spacers between two CRISPR arrays (1 and 2). We modified each CRISPR to only contain a single spacer (1.1 and 2.1), as previously described [19]. The native *cas3* and *cas8* promoters were replaced with *tac* and *araBp8*, respectively using λ-Red recombinase [50]. Four different designed seed sequences (MS1: 5′-TTAAAATT-3′, MS2: 5′-CCGGGGCC-3′, MS3: 5′-TTAAGCCC-3′, MS4: 5′-CCGGATTT-3′) were introduced into the CRISPR 2 locus using a previously developed method for scar-less λ red chromosomal manipulation based on a kanamycin resistance marker and mPheS (modified *E. coli* phenylalanine tRNA synthetase) for counter selection [51]. Outside of the seed sequence, the CRISPR 2 spacer 1 sequence was retained. Following replacement of the seed sequences, a Δ*cas1* strain was constructed for MS1-MS4 using λ-Red recombination [50].

### Generation of seed library

To generate partially randomized seed sequence libraries, four different single-stranded oligonucleotides with 8% doping frequency were designed and synthesized (Integrated DNA Technologies). Here, each position in the seed sequence has 76% chance of having the correct nucleotide that matches the crRNA [47]. The crRNA of Cascade does not base pair with every sixth nucleotide of the protospacer target strand [20,31,38,45,46]. Hence, we utilized this unique sixth position to serve as another barcode for analyzing the high-throughput sequence data (see below), with each library having a different nucleotide at the sixth position (MS1 = A; MS2 = G; MS3 = C; MS4 = T). Oligonucleotides were assembled with pACYC-GFP [52] using the NEBuilder HiFi DNA Assembly Cloning Kit and the products were transformed into NEB Stable competent cells as per the manufacturer’s protocol. Around 50,000 colonies were generated for each library on LB-agar plates supplemented with 34 µg/mL chloramphenicol. The colonies were resuspended with 50 mL of LB supplemented with chloramphenicol (34 µg/mL) and grown overnight at 37 °C with shaking at 180 rpm. The plasmid libraries were then extracted using a Qiagen Miniprep DNA Purification kit. Plasmid selection and high-throughput sequencing was performed for two biological replicates.

### Plasmid library depletion assay

Each library was transformed into the corresponding WT and Δ*cas1* strains. During the recovery stage, the cells were then directly cultured in 50 mL of LB with chloramphenicol and grown overnight at 37 °C with shaking at 200 rpm. This culture was used as cycle 0. The next day, 20 µL of each culture was inoculated into 2 mL fresh LB supplemented with 2 mM IPTG, 20 mM arabinose and no antibiotics. The culture was grown for 10 h, with time points taken every 2 h, or for five 10-h cycles, with time points taken for each cycle and sub-culturing in fresh media with inducers at the end of each cycle. The genomic DNA from each time point or cycle was extracted for use as templates for PCR. For each time point, new spacer acquisition was detected via PCR of the CRISPR arrays within the genomic DNA using Taq DNA polymerase. Newly acquired spacers in CRISPR 2 were detected via PCR using primers listed in Table S1 that anneal to the leader sequence and the constant region of the first spacer sequence.

### Preparation of samples for high-throughput sequencing

To prepare samples for Illumina sequencing, the original library miniprep and genomic DNA from each timepoint/cycle were used as templates for PCR amplification of the target region using Q5 High Fidelity DNA polymerase (New England Biolabs) with Nextera adapters (Table S1). The 132 bp product was then analyzed by 2% agarose gel electrophoresis and purified by QIAquick PCR Purification Kit (Qiagen). These products were then amplified by PCR using Q5 High Fidelity DNA polymerase (New England Biolabs) using a pair of primers containing unique barcodes to differentiate between libraries and replicates. The 224 bp product was analyzed by 2% agarose gel electrophoresis and purified by QIAquick PCR Purification Kit. The samples were mixed, and equal amounts were pooled based on their absorbance reading. Samples were analyzed using an Agilent 2100 Bioanalyzer to determine the size and purity and submitted to Admera Health (New Jersey) or the Iowa State DNA Facility for Illumina MiSeq analysis with paired-end reads of 75 or 150 cycles.

### High-throughput sequencing analysis

Python scripts used for analysis of plasmid library sequencing data are available at github.com/sashital /typeIE_seed_sequence. The scripts were written in collaboration with ChatGPT (OpenAI, January, 2026 model) and were extensively validated.

Seed sequences were extracted based on matching of the constant sequences upstream and downstream of the seed and the presence of a barcoded sixth position described above. The number of reads for each unique seed sequence for each cycle of a give replicate for a given strain was output as a tab-separated file.

Next, the mismatch frequency (MM frequency) was calculated for each number of MM (*n*) at each cycle (*c)* (Fig. S1A).

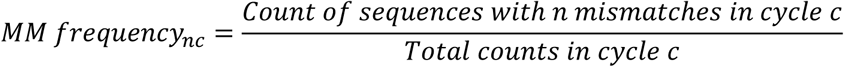

To adjust the abundance of different enriched sequences, a normalization factor (NF) was calculated by averaging the MM frequency of sequences with three, four and five mismatches from each cycle (*c =* 1, 2, 3, 4 or 5), and dividing the average 3-5 MM frequency at cycle 0 by the value at each individual cycle.

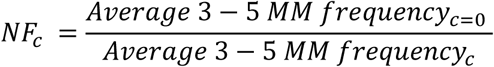

Each NF was multiplied by the frequency from the original MM distribution to generate a normalized MM frequency (Fig. S1B).

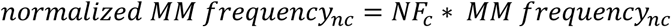

For 1MM and 2MM depletion curves (Fig. 1D-E), the normalized MM frequency for each cycle was divided by the MM frequency at cycle 0 prior to plotting versus cycle number or time point.

The frequency of individual 1MM or 2MM target sequence was measured by dividing the number of counts for sequences with the specific number of mismatches by the total count at a given 10-h cycle or 2-h time point.

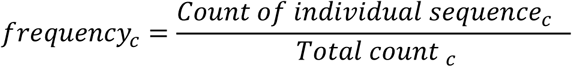

The frequency relative to the control was determined by dividing the frequency at a given cycle by the frequency of the same sequence at cycle 0.

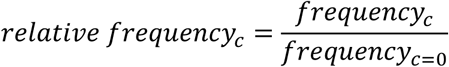

For data that were fit to a delayed exponential decay model (see below), the original library miniprep was used as the control, as we observed a small amount of target loss during cycle 0, likely due to leaky expression of the *cas* operon.

The normalized frequency was determined by multiplying the relative frequency for a given sequence at a given cycle by the normalization factor for that cycle.

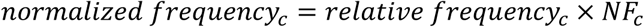

To determine the half-lives show in Fig. 2A, normalized frequency values for 2-hour time points were fit using a delayed exponential decay model in GraphPad Prism 10 (individual plots are shown in Fig. S3). The model assumes an initial plateau followed by first-order decay:

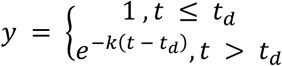

where *y* is the normalized frequency at a given time point *t*, *k* is the decay rate constant, and *t*_d_ is the delay time before decay begins. The initial value was fixed at 1 and the decay plateau was fixed at 0. The half-life was determined by adding the delay time to the half-life calculated using the decay rate constant:

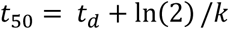

Fits were performed independently for each replicate for both the wild-type and *cas1* deletion strain. Sequences that did not decay sufficiently to obtain a good fit were capped at a half-life of 20 h.

For the comparison of 1MM sequence depletion between the WT and *cas1* deletion strain across 10-h cycles in Fig. 2B, the normalized frequency of each 1MM sequences at each cycle from the WT and Δ*cas1* strains was calculated and the difference was assessed by the depletion score (Z).

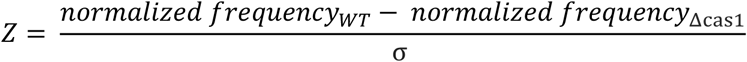

where σ is the standard deviation of all 1MM sequences in the WT and Δ*cas1* strain for a given replicate at a given cycle. Negative values indicate the depletion of the 1MM sequences in the WT but not Δ*cas1* and values close to 0 represent sequences that were depleted in both WT and Δ*cas1*.

For the heatmaps in Fig. 3, the relative frequency of all 2MM sequences were plotted for cycles 1-5. In the development of the heatmaps, we plotted both relative frequency and normalized frequency. The relative frequency plots without normalization to the 3-5MM sequences provided the best contrast between sequences that are tolerated and sequences that cause defects in interference and/or priming and are used in Fig. 3. The average of two replicates is plotted for each sequence at each cycle.

### Plasmids used for protein expression and target cleavage

All plasmids used in this study are listed in Table S3 and were verified by Sanger sequencing and whole plasmid sequencing (Eurofins Genomics). Plasmids used for expression of Cas3 and wild-type or minimal cysteine variants of Cascade without the Cas8 subunit (hereafter Cascade_-8_) and Cas8 are previously described [2,19,39]. For expression of Cascade bearing the MS1 spacer, a CRISPR array containing 8 repeats and 7 MS1 spacers in pET-52b(+) was synthesized by GeneScript and subcloned into pRSF under control of a T7 promoter with *lac* operator by Gibson Assembly. MS1 target plasmids were constructed using pUC19. The target sequences (Table S1) were inserted between BamHI and EcoRI cut sites using restriction cloning. Target plasmids were purified using Qiagen Plasmid Midi Kit.

### Protein purification

All proteins were expressed in BL21(DE3) cells. Overnight cultures were used to inoculate large-scale cultures for protein expression. All cultures were grown to 0.5 OD600 at 37 °C and induced overnight at 20 °C with 0.5 mM isopropyl β-D-1 thiogalactopyranoside (IPTG).

For wild-type Cascade_-8_, cells were lysed in Lysis Buffer 1 (50 mM sodium phosphate dibasic pH 8.0, 500 mM sodium chloride, 5% glycerol, 10 mM imidazole pH 8.0, cOmplete™, EDTA-free Protease Inhibitor Cocktail, 1 mM dithiothreitol (DTT)) using an Avestin homogenizer. Cell debris was removed by centrifuging the cell lysate at 19,000 rpm for 30 minutes. Supernatant was collected and run through HisPur Ni-NTA affinity resin in Lysis Buffer 1, washed with Wash Buffer 1 (50 mM sodium phosphate dibasic pH 8.0, 500 mM sodium chloride, 5% glycerol, 25 mM imidazole pH 8.0, 1 mM DTT) and eluted with Elution Buffer (50 mM Sodium phosphate dibasic pH 8.0, 500 mM sodium chloride, 5% glycerol, 250 mM imidazole pH 8.0, 1 mM DTT). The eluted proteins were cleaved by tobacco etch virus (TEV) protease overnight at 4 °C to remove the His_6_-tag while dialyzing against Dialysis Buffer (50 mM Sodium phosphate dibasic pH 8.0, 500 mM Sodium chloride, 5% glycerol, 2 mM DTT). The cleaved proteins were flowed through a Ni-NTA column to remove uncleaved protein, concentrated to 1 mL, and purified on a Superdex 200 column in Size Exclusion Buffer 1 (20 mM Tris pH 7.5, 100 mM NaCl, 5% glycerol and 1 mM DTT). Minimal cysteine Cascade_-8_ with K169C Cas5 (hereafter K169C-Cascade_-8_) was purified in the same way, except using Tris(2-carboxylethyl)phosphine (TCEP) instead of DTT.

Cas3 was purified as above with the following differences in buffers. Cells were lysed in the Lysis Buffer 2 (20 mM Tris-HCl pH 8.0, 100 mM sodium chloride, 1% glycerol, 10 mM imidazole pH 8.0, cOmplete™, EDTA-free Protease Inhibitor Cocktail, 2 mM DTT) using an Avestin homogenizer. Supernatant after centrifugation was run through HisPur Ni-NTA affinity resin in Lysis Buffer 2, washed with Wash Buffer 2 (1 M NaCl, 25 mM imidazole pH 8.0, 10% glycerol, 2 mM DTT) and then washed with Wash Buffer 1 and eluted with Elution Buffer. Size exclusion was performed in Size Exclusion Buffer 2 (20mM Tris-HCl pH 8.0, 200 mM NaCl, 5% glycerol, 2 mM DTT).

Wild-type Cas8 was purified via HisPur Ni-NTA affinity resin as described above for Cascade_-8_. The eluted proteins were cleaved with TEV protease overnight at 4 °C to remove the MBP-His_6_-tag while dialyzing against Buffer A (50 mM HEPES pH 7.0, 50 mM Sodium chloride, 5% glycerol, 2 mM DTT). Cleaved samples were applied to HiTrap HP-SP (GE Healthcare Life Sciences) ion exchange column equilibrated with Buffer A to separate the MBP-His_6_ tag from Cas8. The column was washed with 10% Buffer B (50 mM HEPES, pH 7.0, 1 M NaCl, 5% glycerol, and 1 mM DTT). Bound proteins were eluted by a gradient from 10% to 50% Buffer B. The sample was concentrated to 1 mL and applied to Superdex 200 column in Size Exclusion Buffer 1 (20 mM Tris pH 7.5, 100 mM NaCl, 5% glycerol and 1 mM DTT). Minimal cysteine H69C Cas8 (hereafter H69C-Cas8) and minimal cysteine N376C Cas8 (hereafter N376C-Cas8) were purified in the same way, except using TCEP instead of DTT.

### Protein labeling for FRET Assays

Cy3-maleimide and Cy5-maleimide (Lumiprobe) were dissolved in 50% DMSO to 4 mM. All labeling reactions were performed in the labeling/cleavage buffer (50 mM HEPES NaOH pH 7.5, 100 mM KCl, 5% glycerol, and 1 mM TCEP). H69C-Cas8 was used to label the N-terminal domain of Cas8 and N376C-Cas8 was used to label the C-terminal domain of Cas8. Each Cas8 variant was labeled with the Cy3 donor fluorophore, and minimal cysteine Cascade_-8_ was labeled with the Cy5 acceptor fluorophore. 20 μM of a Cas8 variant or K169C-Cascade_-8_ was mixed with 200 μM Cy3 or 200 μM Cy5, respectively, with a final DMSO concentration of 5%. Reactions were incubated in the dark at 4 °C for 1 h. Reactions were quenched by adding 10 mM DTT and labeled proteins were separated from free dye using a Spin-X UF 10K MWCO concentrator (Corning). For each set of experiments, proteins were labeled just prior to the experiment.

### DNA substrates for FRET assays

DNA substrates for FRET assays (Table S1) were chemically synthesized by Integrated DNA Technologies and purified by denaturing gel electrophoresis using 12% polyacrylamide with 8 M urea in 1X tris-boric acid-EDTA buffer. DNA was excised from the gel and recovered by crushing the gel pieces and soaking in ddH_2_O overnight at 4 °C. The gel pieces were removed by filtration using Costar Spin-X centrifuge tube filters (Corning), and ethanol precipitated. The dried pellet was resuspended in ddH_2_O and concentration was determined using A260 measured on a nanodrop (Thermo Fisher Scientific). The non-target strand was annealed with the target strand at a 1.2:1 molar ratio in labeling/cleavage buffer by heating at 95 °C for 5 min and cooling to room temperature. To ensure complete binding, the non-target strand was truncated 10-nt after the PAM sequence, as previously described [39].

### FRET sample preparation, measurement, and analysis

All fluorescence measurements were conducted in labeling buffer. 150 nM each Cas8-Cy3 variant and 100 nM K169C-Cascade_-8_ -Cy5 were mixed in 100 μL reactions and incubated at room temperature for 30 min to form the Cascade complex. Cascade complex was mixed with the DNA target at a 1:1.5 ratio and incubated at room temperature for 30 min. For each sample, control experiments were performed to detect donor or acceptor fluorescence in the absence of their corresponding FRET pair. These samples were prepared identically to FRET samples, but unlabeled K169C-Cascade_-8_ (donor alone) or Cas8 variant (acceptor alone) was substituted to enable DNA binding but prevent FRET.

After incubation, the fluorescence of reactions was detected at 25 °C using black 96-well plates (Thermo Fisher Scientific) and a Tecan Spark Multimode Microplate reader. The protocol was set to 10 flashes with 100% gain, Z-position was set to 20000 μm, settle time was set to 5 ms with 10 μs lag time and 50 μs integration time. Excitation filter was set to 485(20) nm and emission wavelength measured from 553 nm to 800 nm with 5 nm bandwidth with 2 nm increments. Fluorescence measurements were performed in triplicate. Fluorescence intensity spectra shown in Fig. 4C-D were normalized and smoothed using GraphPad Prism. The raw fluorescence intensity spectra, including controls, are shown in Fig. S4.

Raw data was analyzed using a previously described method [39]. *E_FRET_* was determined by calculating donor fluorescence quenching [53]:

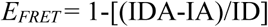

where IDA is the donor fluorescence intensity in the presence of the acceptor, IA is the acceptor fluorescence in the absence of donor, and ID is the donor fluorescence intensity in the absence of the acceptor. All values were calculated by integrating the area under the peaks from 550 to 620 nm, using GraphPad Prism. The change in *E_FRET_* (Δ *E_FRET_*) was calculated by subtracting the *E_FRET_* value for apo Cascade from Cascade bound to targets. The average of three replicates was determined, and standard deviation was propagated for subtraction to determine error.

### Plasmid degradation assays

All reactions were performed in labeling buffer, with the addition of 10 μM CoCl_2_, 10 mM MgCl_2_ and 2 mM ATP (hereafter cleavage buffer). Reactions were performed at room temperature to allow measurement of degradation rates for the perfect target, which occurs too rapidly to perform kinetic analysis at 37 °C. Plasmid degradation was performed through two different types of assays. For the experiments in which Cascade was not pre-bound to DNA, 150 nM wild-type Cas8, 100 nM wild-type Cascade_-8_ and 500 nM Cas3 were pre-mixed. Then, 6 nM target plasmid was added, and the reaction was incubated at room temperature. At each time point, an aliquot was removed, and the reaction was terminated by addition of phenol-chloroform in 1:1 ratio. For experiments in which Cascade was pre-bound to DNA, to eliminate the effects of binding defects on the rate of target degradation, 150 nM wild-type Cas8 and 100 nM wild-type Cascade_-8_ were mixed with 6 nM plasmid. The binding reactions were incubated for 30 min at room temperature. Then, 500 nM Cas3 was added, and aliquots of the reaction were terminated by addition of phenol-chloroform for each time point in 1:1 ratio. For both experiments, the aqueous layer was extracted and mixed with a 2X gel loading solution in 1:1 ratio and analyzed by electrophoresis on a 1% agarose gel with post-staining using SYBR Safe Gel Stain (Thermo Fisher Scientific).

The intensity of DNA bands was quantified by densitometry using ImageJ software. Cas3 initially nicks the DNA followed by degradation, resulting in an initial slow migrating relaxed band and eventual linearization and smearing of the DNA throughout the lane (Fig. S5). The intensities of the nicked, linear, negatively supercoiled, and degraded DNA were measured separately. To calculate fraction cleaved, the product intensity (including nicked, linearized, and degraded DNA) was divided by the total DNA (including supercoiled, nicked, linearized, and degraded DNA). For conditions where a nicked band appeared prior to cleavage initiation, the intensity of the nicked DNA at time point 0 was subtracted from the intensity of the nicked DNA at a given time point prior to determining fraction cleaved. Each measurement was performed in triplicate.

For fitting fraction cleaved versus time, most of the rate curves fit best to a double-exponential rate equation

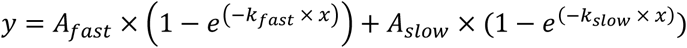

where *A*_fast_ and *A*_slow_ are the amplitudes of the fast and slow phase, respectively, and *k*_fast_ and *k*_slow_ are rate constants for the fast and slow phase, respectively. All rate curves were fit to this equation with the exception of the perfect target (rU-dA) for both conditions and the rU-dC target when Cascade was not pre-bound to the target, which fit better to a single-exponential rate equation

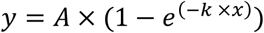

The data were fit in GraphPad Prism. The fits of average data from the three replicates are shown in Fig. 5D.

To summarize rate constants measured using the double-exponential rate equation, *k*_ave_ was calculated by using the following formula:

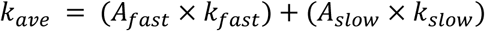

For the targets where a single-exponential rate equation was used, the *k* value determined from the fit is reported. Values for *k* or *k*_ave_ were determined for each replicate, and average values were plotted in Fig. 5E, with error representing the standard deviation of the three replicates.

## Author Contributions

P.P. – Conceptualization, investigation and formal analysis for plasmid library depletion assays, Writing – original draft.

M.O. – Conceptualization, investigation, formal analysis, and visualization for FRET and cleavage assays, Writing – original draft.

E.A.D. – Investigation for FRET assays.

J.R. – Investigation for FRET assays

C.X. – Conceptualization and methodology for construction of the original inducible strain.

D.G.S. – Conceptualization, formal analysis, visualization, supervision, and funding acquisition for project.

## Acknowledgements

The authors thank Michael Baker of the Iowa State University DNA Facility for assistance with Illumina sequencing; Nicholas Crumpton for assistance with the Tecan Spark Microplate reader; and other members of the Sashital lab for helpful discussions and experimental suggestions. This study was supported by a grant from the National Institutes of Health (R35 GM140876).

## Supplementary Information

**Figure S1:**
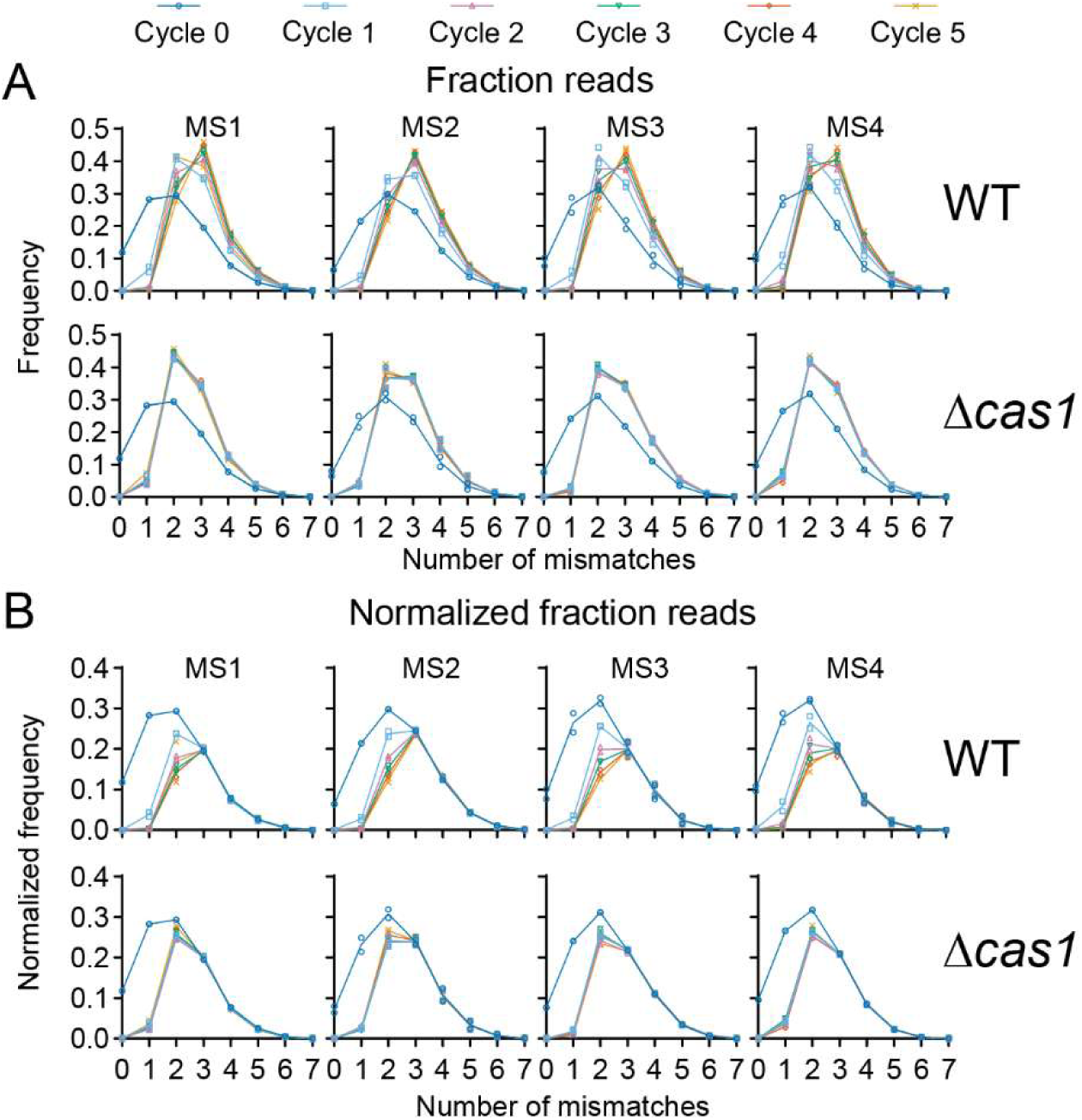
Distribution of sequences with 0 to 7 mismatches in each plasmid library at each cycle. (A) The fraction of reads for sequences containing each number of mismatches is plotted prior to normalization. (B) The fraction of reads was normalized relative to the frequency of sequences containing 3, 4, or 5 mismatches (see Methods). Lines represent the average of two separate replicates. Individual data points for both replicates are plotted.

**Figure S2:**
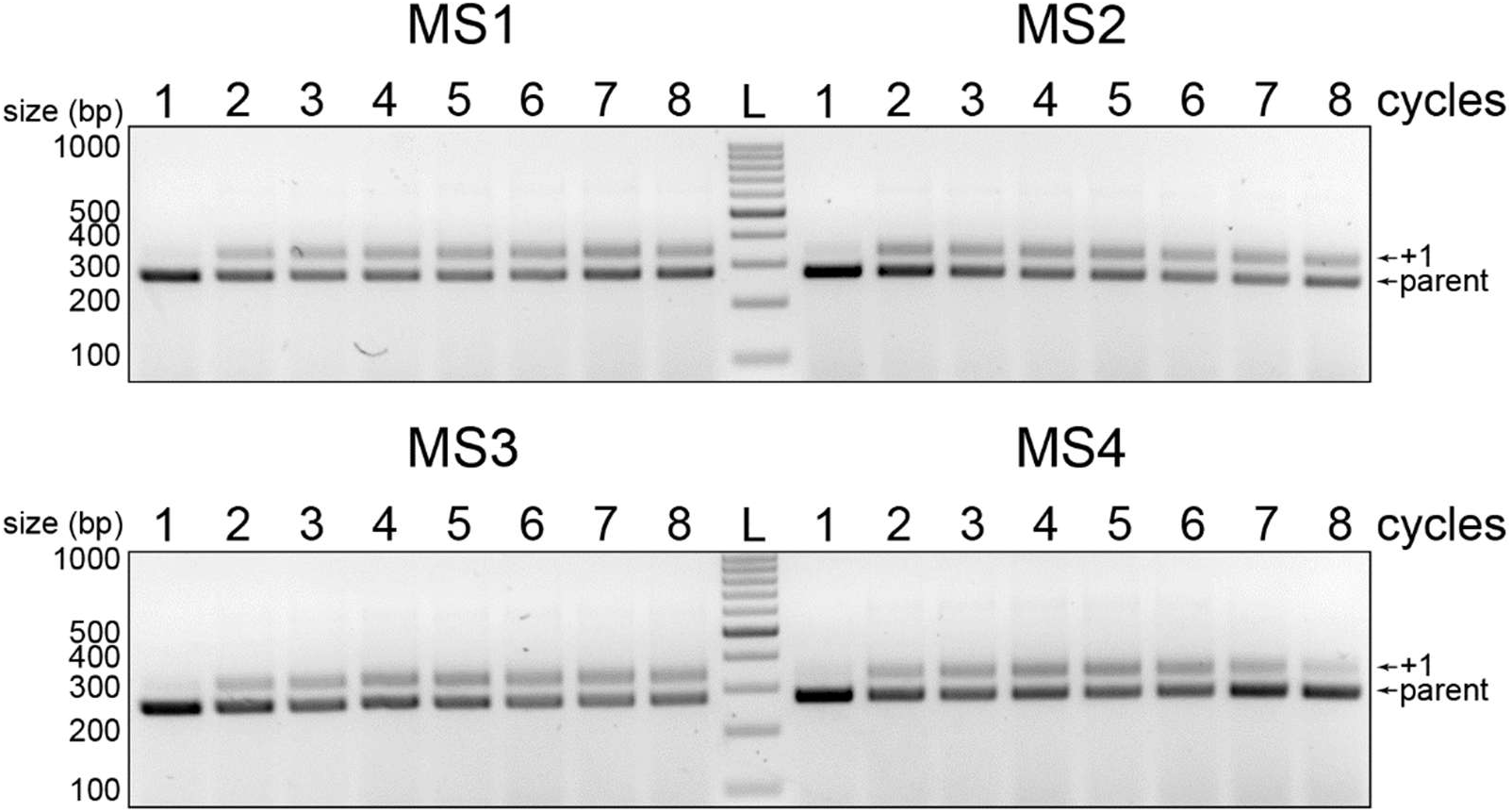
Spacer acquisition across 10-h cycles for MS1-MS4 WT strains. PCR amplification of the CRISPR 2 array was used to detect CRISPR expansion in each strain bearing the corresponding plasmid library. The +1 band appears following the acquisition of new spacers into the CRISPR array. L = 100 bp ladder.

**Figure S3:**
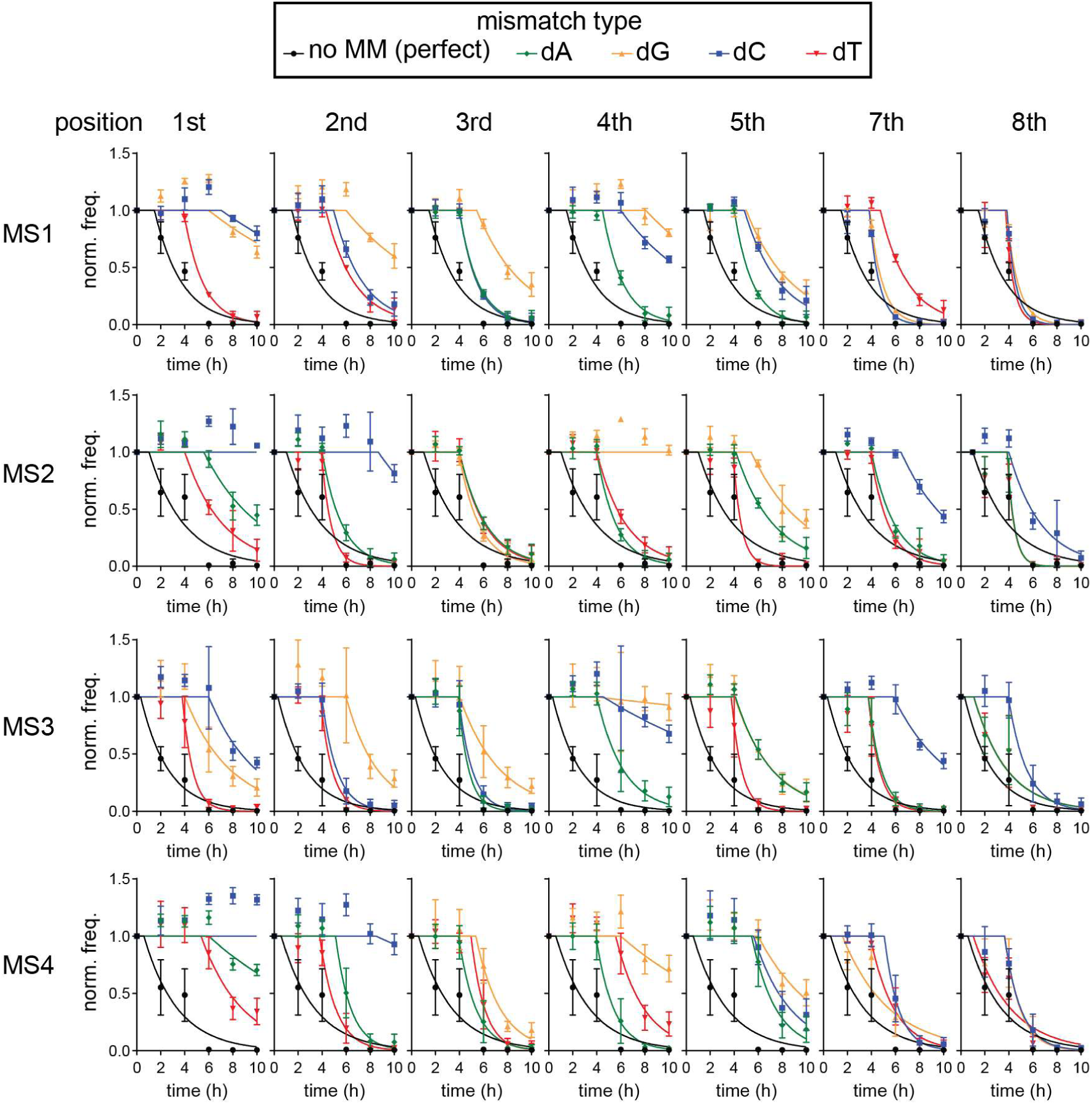
Decay curves for 1MM target sequences of WT strains used to calculate half-lives in Fig. 2. Each graph plots normalized frequency (norm. freq.) versus time for the perfect target (black with circles) or targets containing a single mismatch at a given position of the seed. The three possible mismatch types at each position are plotted, with dA shown in green with diamonds, dG shown in orange with triangles pointing up, dC shown in blue with squares, and dT shown in red with triangles pointing down. The average of two WT and two Δ*cas1* replicates are plotted as the data points, and error bars represent standard deviation. The data were fit to an exponential decay following a delay, as described in the Methods.

**Figure S4:**
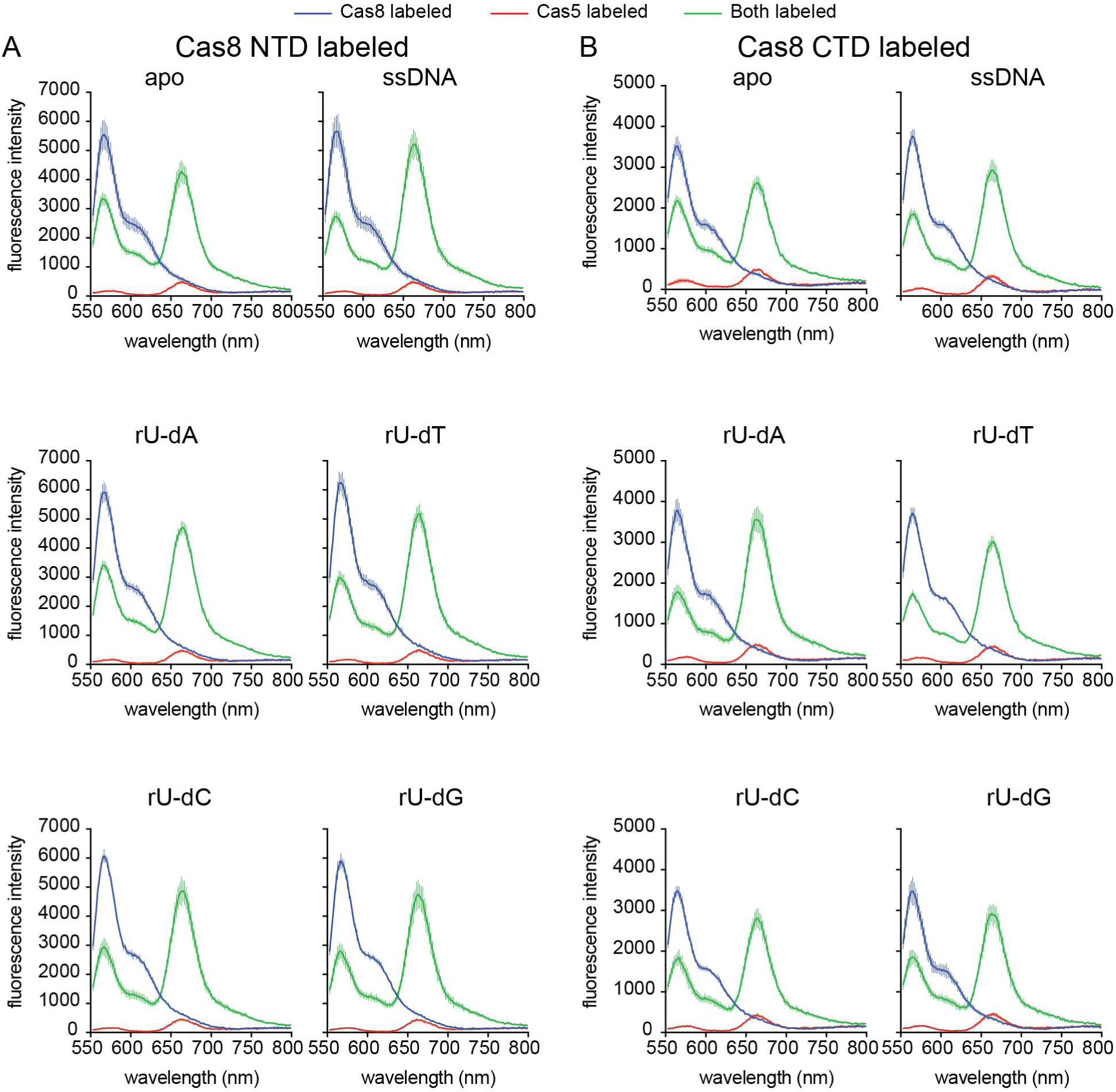
Individual graphs and controls for FRET data. Fluorescence intensity spectra are shown for all samples used to calculate the change in FRET efficiency (Δ*E*_FRET_). The (A) N-terminal or (B) C-terminal domain of Cas8 was labeled. The donor intensity traces in the absence of acceptor (i.e. Cas8-Cy5 with unlabeled Cascade_-8_) are shown in blue, the acceptor intensity traces in the absence of donor (i.e. unlabeled Cas8 with Cy3-labeled Cascade_-8_) are shown in red, and the FRET traces with both fluorophores present are shown in green. For each sample, all three sets of Cascade were bound to the indicated DNA target, with the exception of apo, in which no DNA was added. The average of three replicates is shown, and error bars represent the standard deviation of fluorescence intensity at each wavelength that was measured.

**Figure S5:**
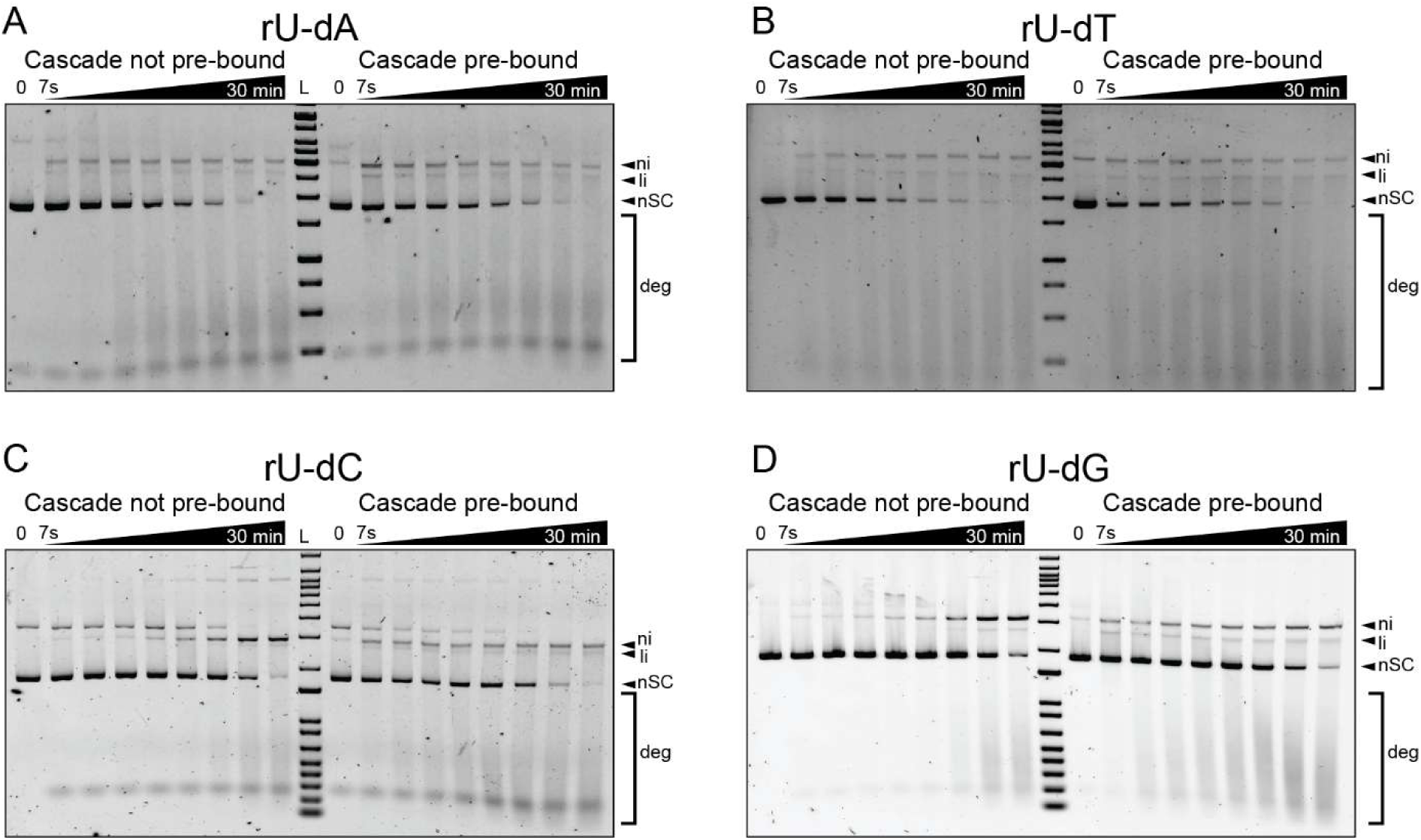
Example gels for cleavage assays. Agarose gels for the cleavage assays that were quantified in Fig. 5D-E. For each gel, a single replicate of a cleavage assay in which Cascade was not pre-bound to the DNA (left) or Cascade was pre-bound to the DNA (right) is shown for (A) the perfect target containing an rU-dA base pair at position 1 or targets introducing an (B) rU-dT, (C) rU-dC, or (D) rU-dG mismatch at position 1. The DNA is initially negatively supercoiled (nSC). Following initiation of cleavage, nicked (ni), linear (li) and degradation (deg) products appear. For conditions where a nicked band appeared prior to cleavage initiation, the intensity of the nicked DNA at time point 0 was subtracted from the intensity of the nicked DNA at a given time point.

**Table S1:**
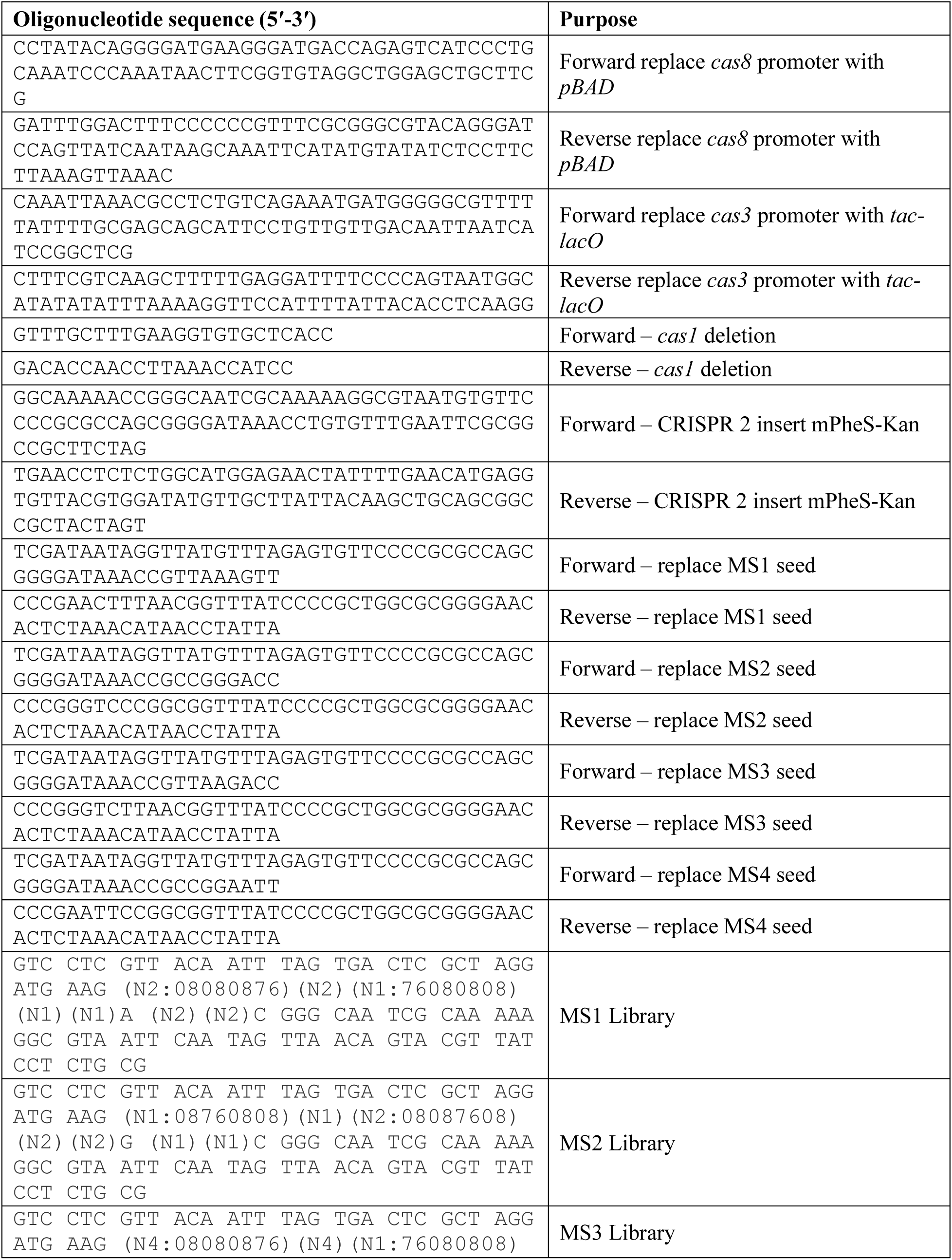
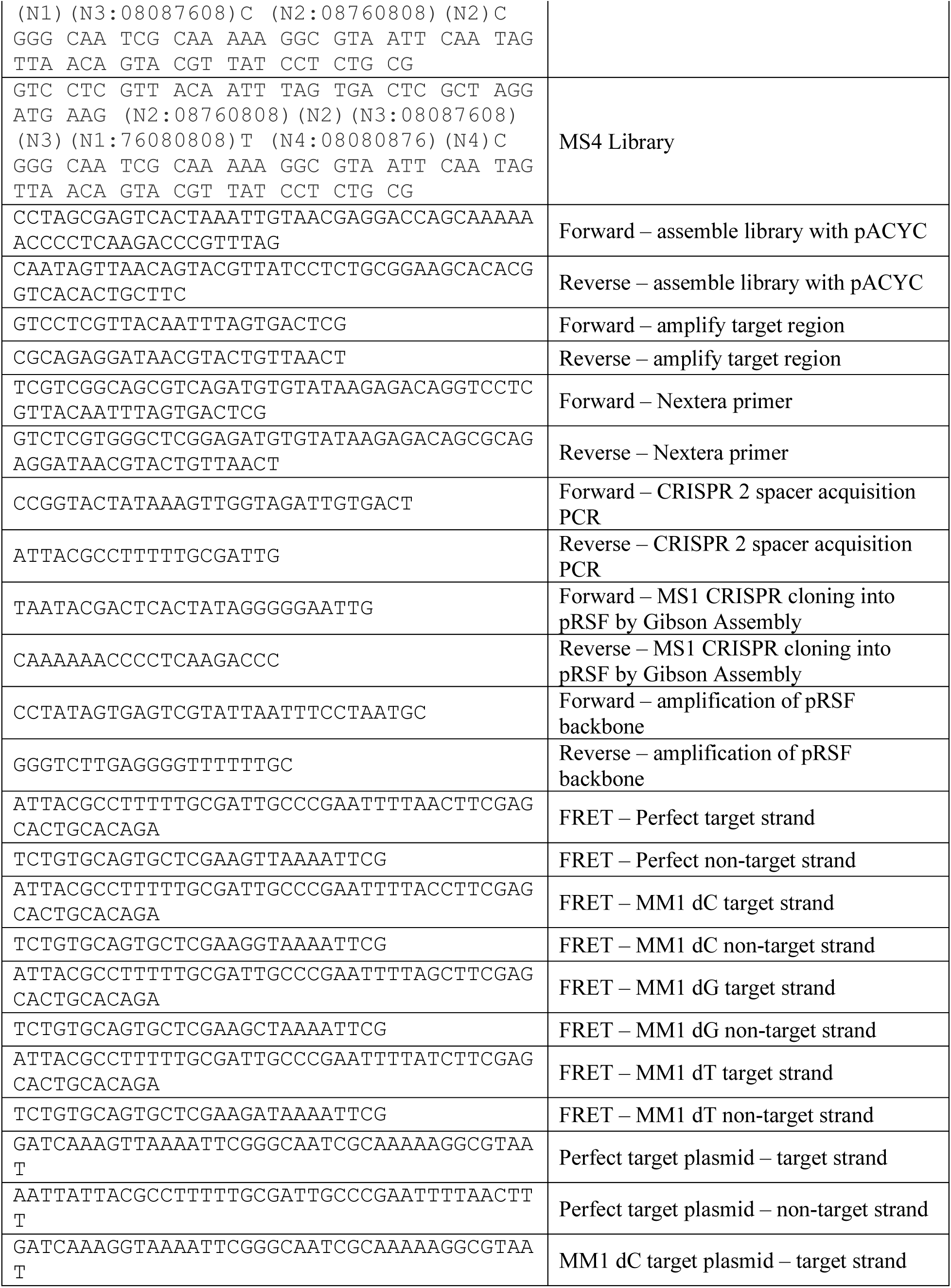
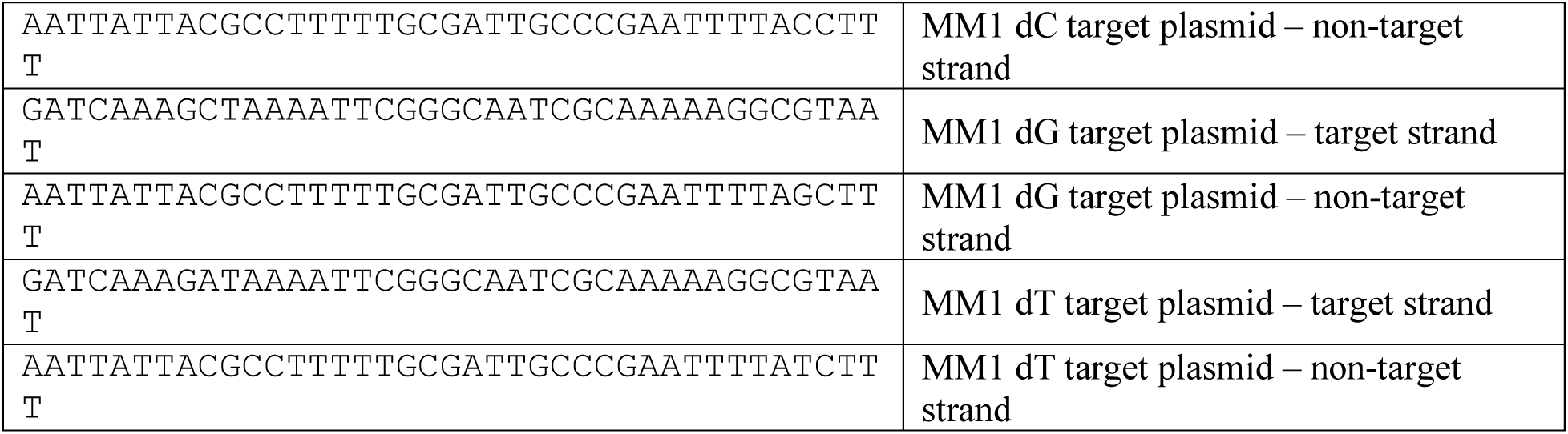
List of oligonucleotides used in this study.

**Table S2:**
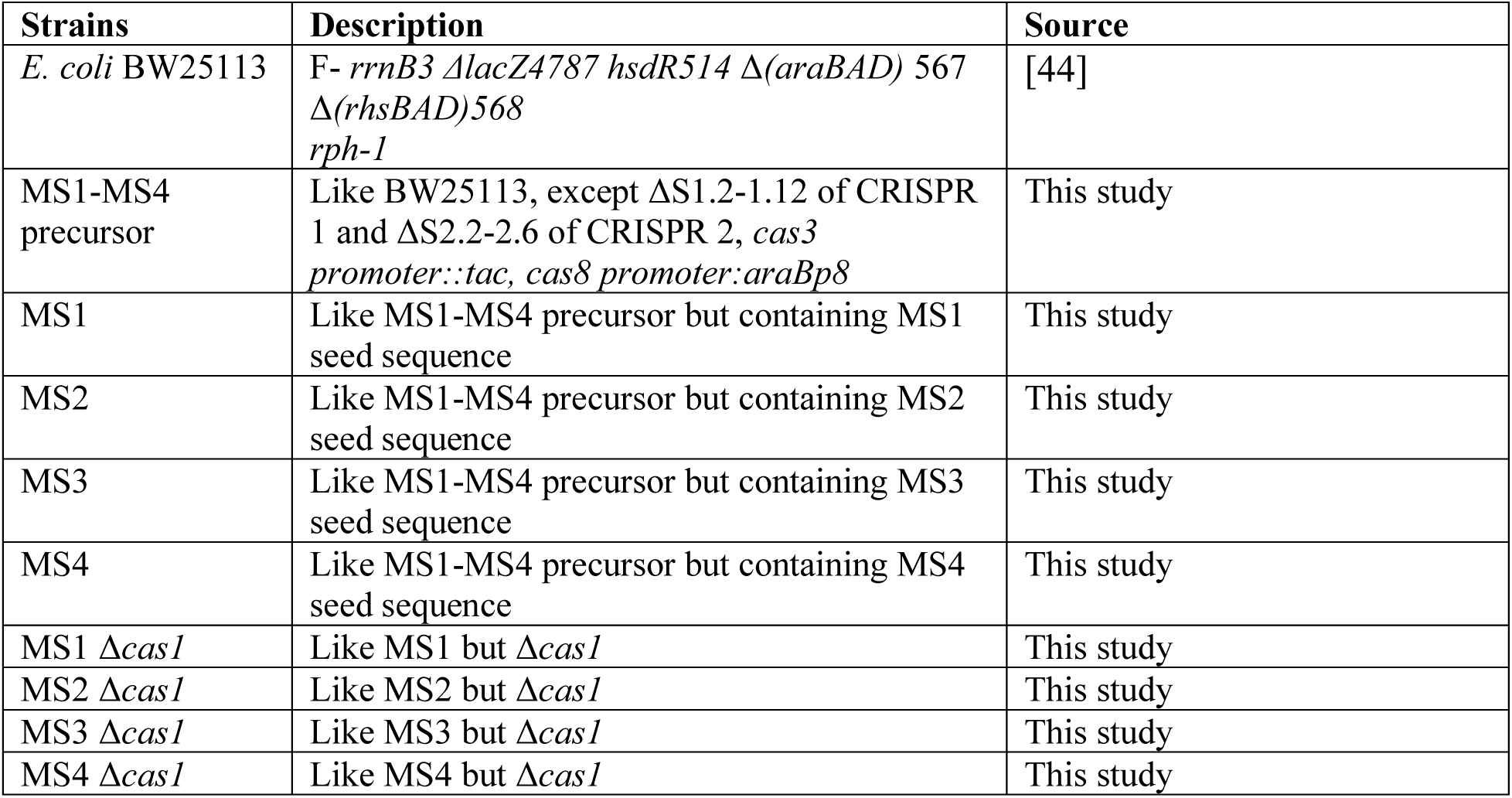
List of strains used in this study.

**Table S3:**
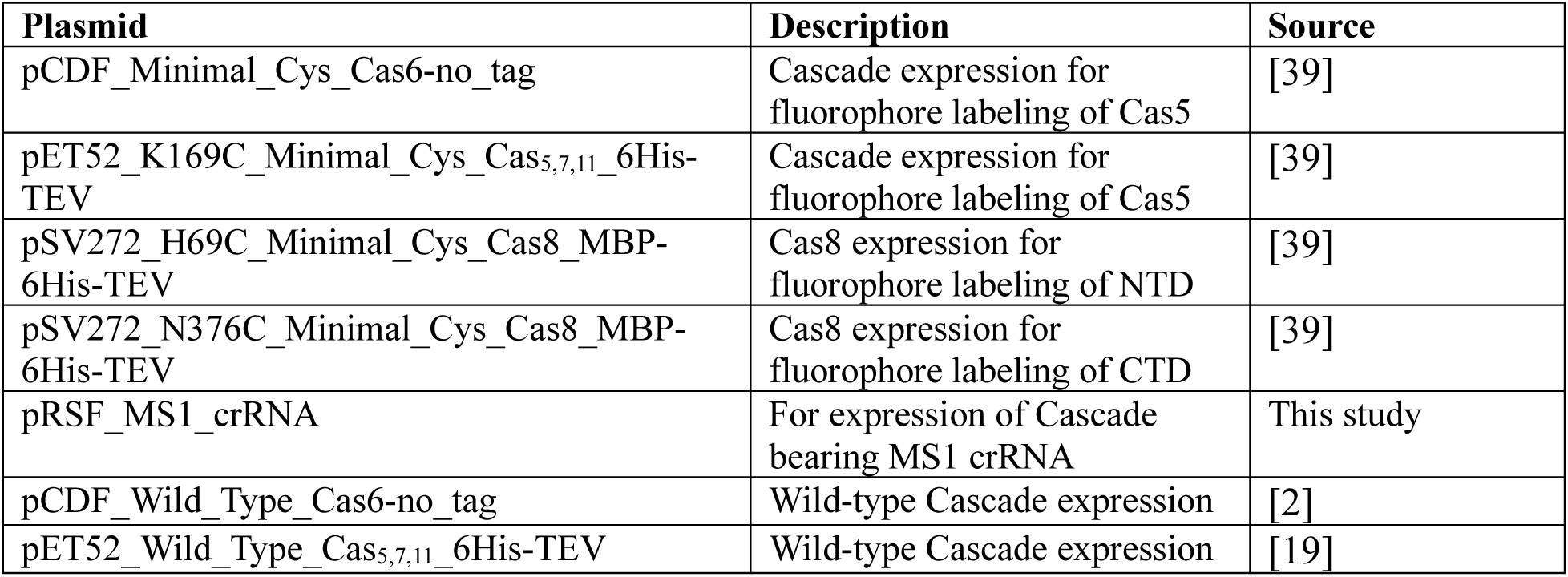
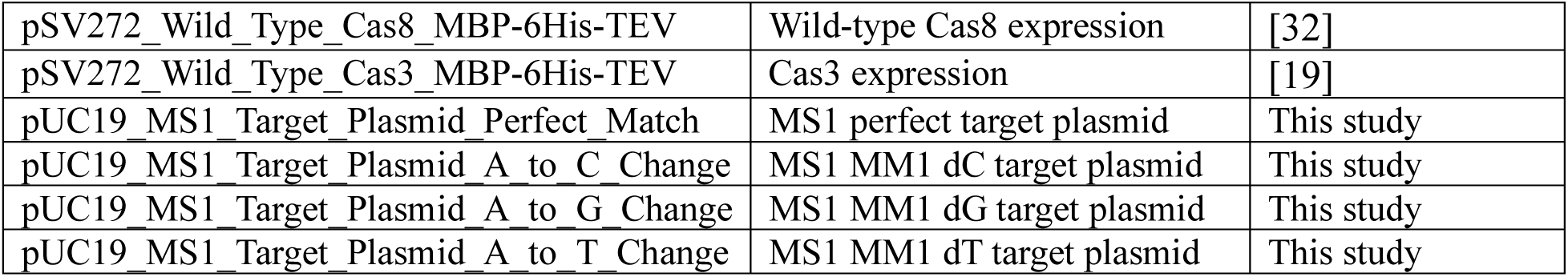
List of plasmids used in this study.

